# Chronic viral infection promotes early germinal center exit of B cells and impaired antibody development

**DOI:** 10.1101/849844

**Authors:** Ryan P. Staupe, Laura A. Vella, Sasikanth Manne, Josephine R. Giles, Wenzhao Meng, Ramin Sedaghat Herati, Omar Khan, Jennifer E. Wu, Amy E. Baxter, Eline T. Luning Prak, E. John Wherry

## Abstract

Chronic viral infections disrupt B cell responses leading to impaired affinity maturation and delayed control of viremia. Previous studies have identified early pre-germinal center (GC) B cell attrition but the impact of chronic infections on B cell fate decisions in the GC remains poorly understood. To address this question, we used single-cell transcriptional profiling of virus-specific GC B cells to test the hypothesis that chronic viral infection disrupted GC B cell fate decisions leading to suboptimal humoral immunity. These studies revealed a critical GC differentiation checkpoint that is disrupted by chronic infection, specifically at the point of dark zone re-entry. During chronic viral infection, virus-specific GC B cells were shunted towards terminal plasma cell (PC) or memory B cell (MBC) fates at the expense of continued participation in the GC. Early GC exit was associated with decreased B cell mutational burden and antibody quality. Persisting antigen and inflammation independently drove facets of dysregulation, with a key role for inflammation in directing premature terminal GC B cell differentiation and GC exit. Thus, these studies define GC defects during chronic viral infection and identify a critical GC checkpoint that is short-circuited, preventing optimal maturation of humoral immunity.

## INTRODUCTION

Effective humoral immune responses to infection and immunization are defined by high-affinity antibodies generated as a result of B cell differentiation and selection that occurs within germinal centers (GC) (Cyster and Allen, 2019; Plotkin, 2008). Within the GC, B cells undergo affinity maturation, an iterative and competitive process wherein B cells mutate their immunoglobulin genes (somatic hypermutation) and undergo clonal selection by competing for T cell help (Mesin et al., 2016; Shlomchik et al., 2019). Balancing the decision to remain within the GC and continue participating in affinity maturation or to exit the GC as a plasma cell (PC) or memory B cell (MBC) is critical for achieving optimal antibody avidity, antibody quantity, and establishing immunological memory in response to immunization or infection.

Much of what is known about GC biology comes from studies of immunization (e.g. with haptens and/or protein antigens) or the analysis of acutely resolved infections. In contrast to immunization or acutely resolved infections, humoral immunity in chronic viral infections is often dysregulated. Hypergammaglobulinemia is a hallmark of chronic viral infections such as human immunodeficiency virus (HIV) (Lane et al., 1983; Milito et al., 2004; Moir and Fauci, 2017), hepatitis C virus (HCV) (Bjøro et al., 1994; Cashman et al., 2014; Racanelli et al., 2006), and during infection with chronic strains of lymphocytic choriomeningitis virus (LCMV) in mice (Hunziker et al., 2003; Oldstone and Dixon, 1969). Despite increased total antibody titers, the antibodies made during chronic viral infections are often of poor quality, displaying lower affinity maturation compared to antibodies detected following acutely resolved infections and delayed development of virus-neutralizing function (Battegay et al., 1993; Gray et al., 2007; Logvinoff et al., 2004). In addition, vaccine-induced antibody responses are of poorer quality and quantity in individuals with pre-existing chronic viral infections compared to healthy control subjects (McKittrick et al., 2013; Shive et al., 2018; Wiedmann et al., 2000). Together, these observations suggest that chronic viral infections may impair B cell differentiation and fate decisions, leading to dysregulated humoral immunity.

Although dysregulated humoral immunity has long been appreciated during chronic viral infections, the key steps in B cell differentiation and underlying mechanisms of the dysregulation are less well understood. Limited access to neutralizing epitopes (Klein and Bjorkman, 2010; Moir and Fauci, 2017; Sommerstein et al., 2015), high viral mutation rates (Bonsignori et al., 2017), establishment of a latent reservoir (Barton et al., 2011), and direct infection of B cells (Barton et al., 2011) are among the immune evasion strategies used by chronic infections to blunt or evade antiviral antibody responses (Virgin et al., 2009). More recent studies have shown a role for type I interferon in driving B cell differentiation away from the GC fate, resulting in the acute depletion of B cells during chronic viral infection before GC B cell differentiation can be established (Fallet et al., 2016; Moseman et al., 2012; Sammicheli et al., 2016). Moreover, chronic viral infections in mice and humans can result in lymphoid tissue disruption and/or fibrosis (Mueller et al., 2007; Schacker et al., 2006; Wilson et al., 2013), perhaps impacting B cell responses. However, despite these challenges, GC still form during chronic infection (Barnett et al., 2016; Daugan et al., 2016) and antibodies have been shown to diversify over time during chronic HIV and HCV infections (Doria-Rose et al., 2014; Kinchen et al., 2018; Wu et al., 2015), suggesting ongoing GC activity. Moreover, despite generally smaller GCs during chronic infection, affinity-matured antibodies produced as a result of the GC response exert partial but critical control of viral replication and, in some cases, lead to viral control (Chen et al., 2018; Chou et al., 2016; Doria-Rose et al., 2014; Kinchen et al., 2018). Additionally, non-neutralizing antibodies are required to partially contain infection in many cases (Barnett et al., 2016; Hangartner et al., 2006; Horwitz et al., 2017; Richter and Oxenius, 2013; Straub et al., 2013) and can apply selective pressure for viral escape and evolution in some settings (Doria-Rose et al., 2014; Kinchen et al., 2018; Wu et al., 2015). Thus, humoral immune responses that occur during chronic infections likely involve considerable participation of GC B cell responses. Yet, the biology governing GC B cell differentiation and fate decisions in the setting of chronic viral infections remains unclear. A better understanding of dysregulated humoral immunity and whether/how GC biology is impacted during chronic viral infections could provide valuable insights to improve vaccination and antibody responses during chronic infections.

To address these questions, we interrogated the impact of chronic viral infection on B cell differentiation and fate decisions within the GC. Focusing on virus-specific B cells, we employed a combination of bulk and single-cell RNA sequencing (scRNA-seq), together with BCR repertoire profiling and *in vivo* cellular experiments. These studies revealed a key GC checkpoint that is dysregulated during chronic infection. Specifically, during chronic viral infection, GC B cells inefficiently re-enter the dark zone (DZ) during the cyclic GC reaction and instead prematurely exit as MBC or terminal PC. This short-circuiting of the cyclic process of GC B cell development occurred at a key LZ step in GC B cell differentiation and was associated with decreased affinity maturation and serum antibody avidity. Single-cell profiling revealed attenuated BCR signaling and amplified inflammatory signals associated with this altered GC developmental pattern. Indeed, although intentional *in vivo* antigen overstimulation increased the magnitude of the B cell response during acute and chronic infection, additional antigen did not further drive skewed B cell differentiation in either infection. Moreover, the attenuation of BCR signaling skewed B cell differentiation towards the PC and MBC fates during acute infection but not during chronic infection. Our studies reveal the roles of the chronic inflammatory environment and antigen burden on B cell differentiation during chronic viral infection and identify key pathways that can be targeted to modulate humoral immune responses and promote affinity maturation during antiviral immune responses and immunization.

## RESULTS

### Chronic infection promotes terminal B cell differentiation

Humoral immunity in chronic viral infections is dysregulated and often associated with poor pathogen control. These defects in B cell responses are characterized by hypergammaglobulinemia, altered B cell differentiation, and the delayed development of neutralizing antibodies (Battegay et al., 1993; Eschli et al., 2007; Gray et al., 2007; Hunziker et al., 2003; Logvinoff et al., 2004; Oldstone and Dixon, 1969). Despite these observations, it has remained challenging to quantify the virus-specific B cell response to chronic viral infections and track numerical changes and differentiation over time. Previous studies in the mouse model of chronic LCMV infection identified a major detrimental effect of excessive type I interferon signaling causing profound B cell attrition at early time points (days 0-3) that led to defects in subsequent humoral immunity (Fallet et al., 2016; Moseman et al., 2012; Sammicheli et al., 2016). Despite these early B cell defects, germinal centers (GC) still form during chronic infections and the biology of GC responses during these persisting infections remains poorly understood (Barnett et al., 2016; Clingan and Matloubian, 2013; Daugan et al., 2016; Fukazawa et al., 2015). To begin to dissect these questions, we interrogated the LCMV-specific B cell response to acutely resolved and chronic viral infection using the LCMV mouse model of chronic viral infection. In this model, the LCMV Armstrong strain (LCMV Armstrong; Arm; Acute) results in an acutely resolved infection that is rapidly controlled by day 8 post-infection (p.i.) whereas LCMV clone 13 (LCMV clone 13; chronic) results in a chronic infection with persistent inflammation and antigen associated with viral persistence in multiple tissues for the life of the host (Wherry et al., 2003). Although the LCMV-GP protein is the target of the neutralizing antibody response, we focused on the LCMV nucleoprotein (NP) response as it is the dominant target of the LCMV-specific antibody response (Battegay et al., 1993). In addition, NP protein sequences are identical between the acutely resolving and chronic strains of LCMV (Matloubian et al., 1990, 1993; Sullivan et al., 2011). We identified LCMV-NP-specific B cells using fluorescently-labeled bacterially-produced LCMV nucleoprotein (NP) (**Figure 1A** and **S1A**) as previously described (Fallet et al., 2016; Schweier et al., 2019; Sommerstein et al., 2015). Acutely resolved or chronic LCMV infection resulted in similar frequencies of LCMV-specific B cells during the first 7-10 days of infection (**Figure 1B**). However, by day 15 p.i., LCMV-specific B cells increased in frequency and total number following acutely resolved infection but remained lower in magnitude during chronic infection (**Figure S1B**). These data indicate that chronic infection results in a decreased frequency of virus-specific B cells during the timepoints when the GC and memory B (MBC) cell responses are developing.

**Figure 1:**
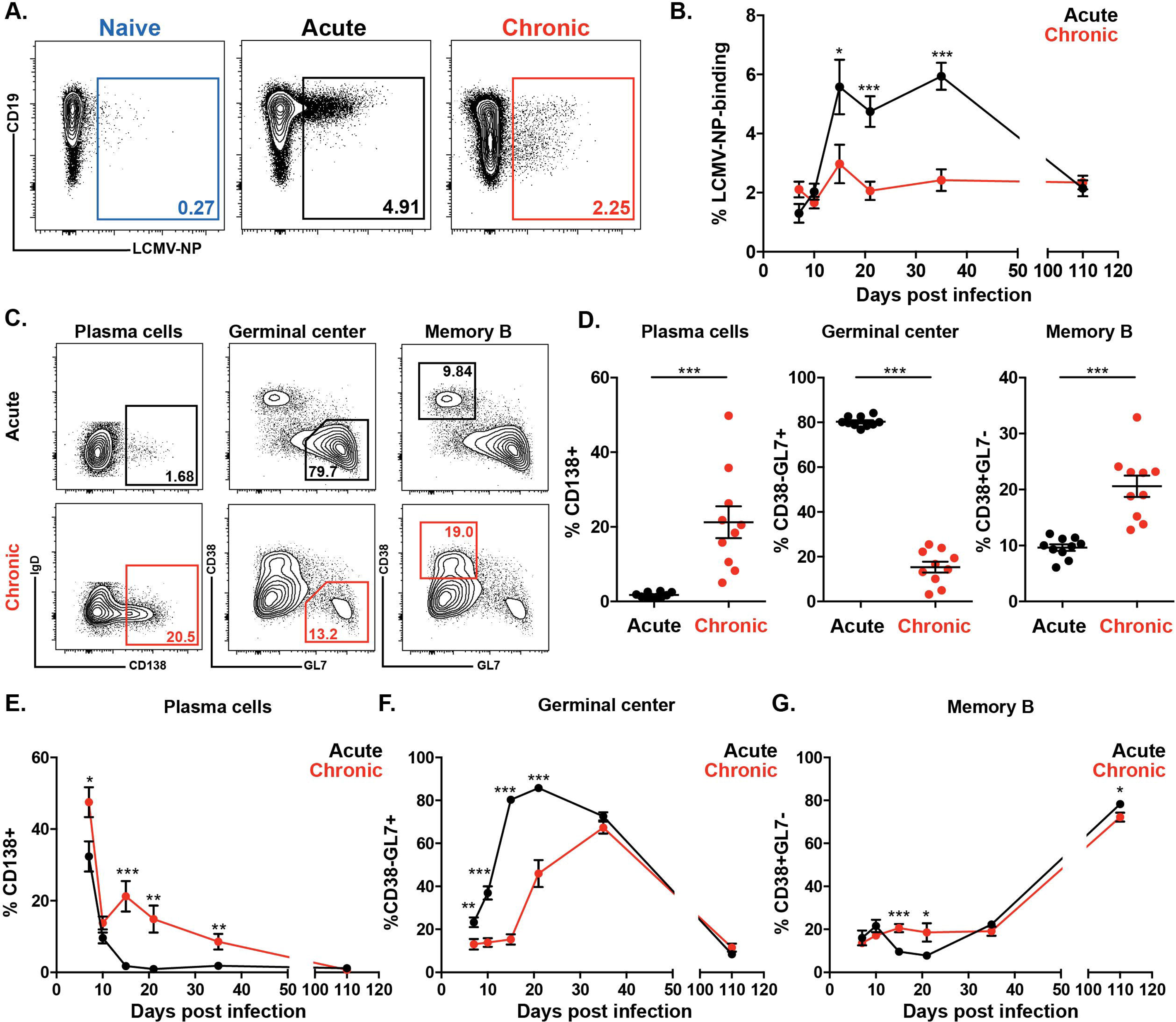
Chronic infection promotes terminal B cell differentiation. **(A)** Representative FACS plots of splenic LCMV-specific B cell staining in naïve, LCMV Armstrong (Acute), and LCMV clone 13 (Chronic) infected mice at d15 post-infection. Cells were previously gated as IgD-B220+Dump-. Dump gate includes CD4, CD8, F4/80, and NK1.1. **(B)** Frequency of splenic LCMV-specific B cells in LCMV Armstrong and LCMV clone 13 infected mice at days 7, 10, 15, 21, 35, and 110 post-infection. **(C)** Representative FACS plots of splenic LCMV-specific plasma cells, germinal center B cells, and memory B cells in LCMV Armstrong and LCMV clone 13 infected mice at d15 post-infection. Cells were previously gated on LCMV-specific B cells as described in A. **(D)** Frequency of splenic plasma cells, germinal center B cells, and memory B cells in LCMV Armstrong and LCMV clone 13 infected mice at day 15 post-infection. **(E-G)** Kinetics of LCMV-specific splenic plasma cell **(E)**, germinal center B cell **(F)**, and memory B cell **(G)** frequencies in LCMV Armstrong and LCMV clone 13 at days 7, 10, 15, 21, 35, and 110 post-infection. Each data point in B, E-G and H-J shows mean ± SEM with 3-5 mice per time point from two independent experiments. Data points in D show mean ± SEM from 10 mice pooled from two independent experiments. *p<0.05, **p<0.01, ***p<0.001, Student’s t-test.

Tight regulation of B cell differentiation into GC B cells, plasma cells (PC) and memory B cells (MBC), is critical for achieving optimal antibody avidity, antibody quantity, and immunological memory (Cyster and Allen, 2019). Thus, to define the LCMV-specific B cell response in more detail, we used markers to distinguish PC (CD138+), GC B cells (CD38-GL7+), and MBC (CD38+GL7-) at day 15 p.i. (**Figures 1C** and **S1A**). In acute infection, approximately 80% of the LCMV-specific B cells were GC phenotype at this time point (**Figure 1C-D**). In contrast, during chronic infection, there were four-fold fewer LCMV-specific GC B cells whereas PC and MBC frequencies were increased 10-fold and two-fold, respectively (**Figure 1C-D**). In chronic infection there was also a population of CD38-GL7-LCMV-specific B cells that was comprised of primarily PC but also a low frequency of age-associated B cells with high expression of Tbet (**Figure S1C**). The reduced frequency of GC B cells was not due to an absence of GCs during chronic infection and the GCs that were present were similar in size to those observed following acutely resolved infection (**Figure S2D**). Thus, at the peak of the virus-specific B cell response, the distribution of LCMV-specific B cell responses during chronic infection was skewed towards the PC and MBC fates and away from the GC B cell fate.

Longitudinal analysis of these LCMV-specific B cell responses during acutely resolved or chronic infection revealed differences in the magnitude and kinetics of PC, GC B cell and MBC responses (**Figure 1E-G** and **S1D-F**). Whereas PC and MBC responses were increased in frequency during chronic infection, especially during the second and third week of infection, GC B cell frequencies were substantially lower during this time frame. These data reveal a sustained quantitative defect in the overall virus-specific B cell response during chronic infection, but also identify differences in PC, MBC, and GC responses over time. Collectively, these data suggest that chronic infection promotes early induction of PC and MBC responses at the expense of the GC response.

### Affinity maturation is impaired during chronic infection

Germinal centers are the primary location of affinity maturation, a cyclic process by which B cells mutate their immunoglobulin genes, compete for antigen and T cell help, and undergo clonal expansion following positive selection (Cyster and Allen, 2019; Mesin et al., 2016). Previous studies have shown that accumulation of affinity-enhancing mutations is critical for eventual control of viremia during chronic LCMV infection (Chen et al., 2018; Chou et al., 2016). However, the delayed production of neutralizing antibody during chronic LCMV infection (Battegay et al., 1993; Eschli et al., 2007) suggests that the process of affinity maturation is dysregulated, an alteration that may also occur during other chronic infections (Doria-Rose et al., 2014; Kinchen et al., 2018; Minamitani et al., 2015; Wu et al., 2015). Based on the data above showing reduced numbers of LCMV-specific GC B cells during chronic infection but increased numbers of LCMV-specific PC, we hypothesized that chronic infection might increase the overall quantity of LCMV-specific antibodies but impair affinity maturation and therefore serum antibody avidity. We first measured serum LCMV-specific antibody titers using an LCMV-NP-specific ELISA. Consistent with our previous longitudinal flow cytometry analysis of LCMV-specific PC, LCMV-NP-specific IgG titers were increased during chronic infection at day 7 p.i. (prior to initiation of the GC response) and at all timepoints measured after day 21 p.i. (**Figure 2A**). We next sought to determine whether LCMV-specific serum antibody avidity was impacted by chronic infection. Thus, we measured LCMV-specific serum antibody avidity using an ELISA combined with a chaotropic sodium thiocyanate (NaSCN) wash to remove low avidity antibody (Nawa, 1992; Pullen et al., 1986). In this assay, antibody avidity (Avidity Index) was measured by comparing the fraction of serum antibody sensitive to removal by NaSCN treatment to total antibody levels (**Figure S2A**). Following acutely resolved infection, the avidity index increased with time p.i., indicating affinity maturation and development of high avidity antibody (**Figures 2B** and **2C**). Serum antibody avidity increased following the formation of the LCMV-specific GC response following acute infection, which started at day 10 and plateaued at day 45 p.i. In contrast to acutely resolved infection, the development of high avidity antibodies was delayed in chronic infection. The avidity indices of the LCMV-specific antibody response were similar between acutely resolving and chronic infection at early time points p.i. but began to diverge at day 35 p.i., with lower avidity antibodies present during chronic infection. Thus, chronic infection impairs and/or delays the development of high avidity antibodies.

**Figure 2:**
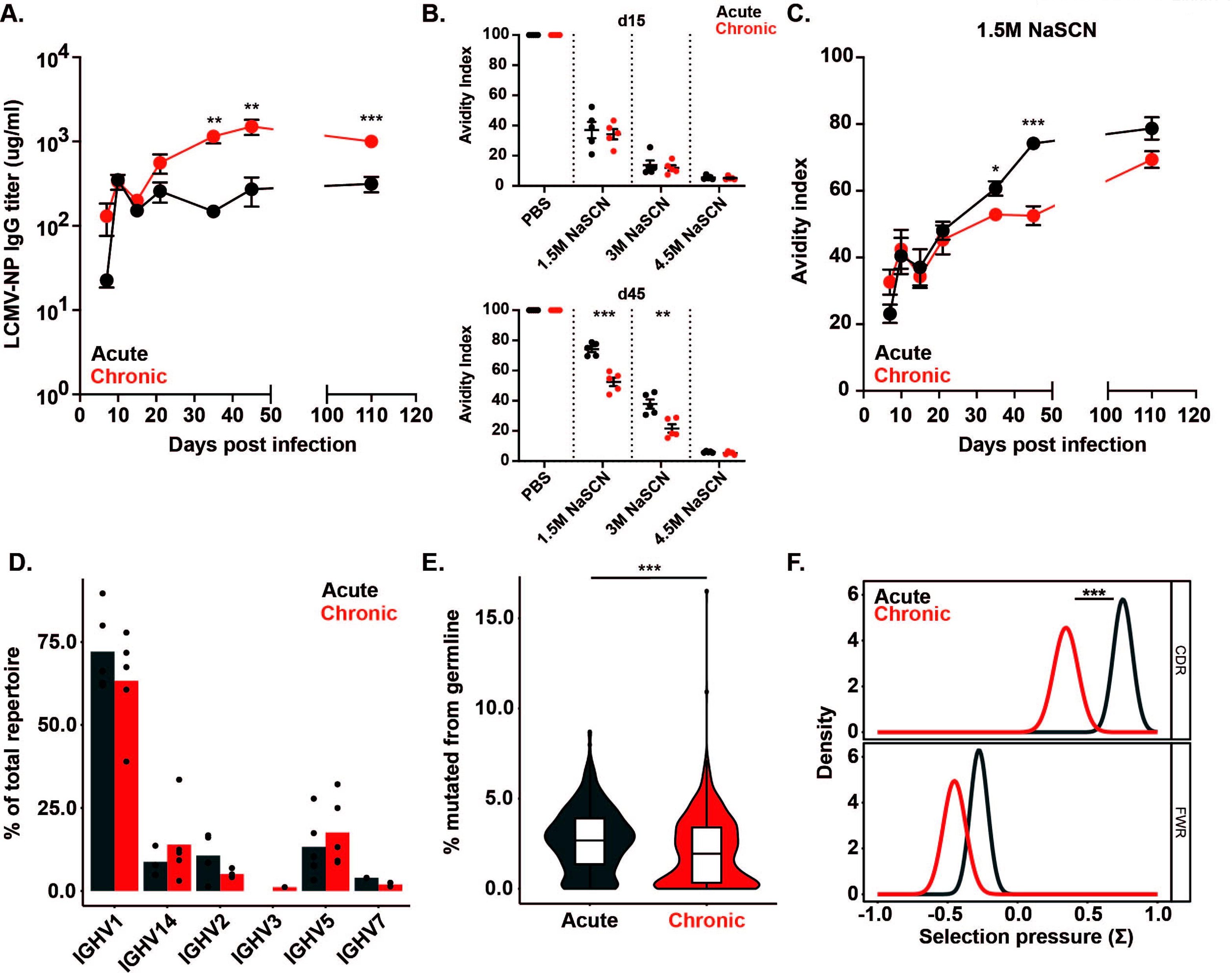
Affinity maturation is impaired during chronic infection. **(A)** Kinetic measurement of serum LCMV-NP-specific IgG titers in LCMV Armstrong (acute) and LCMV-clone 13 (chronic) infected mice at days 7, 10, 15, 21, 35, and 110 post-infection. **(B)** Anti-LCMV-NP serum avidity index measurements in LCMV Armstrong and LCMV clone 13 infected mice at days 15 and 45 post-infection using chaotropic NaSCN ELISA. Dose response to increasing amounts of the chaotropic agent NaSCN is depicted. **(C)** Anti-LCMV-NP serum avidity index in response to 1.5M NaSCN in LCMV Armstrong and LCMV clone 13 infected mice at days 7, 10, 15, 21, 35, and 110 post-infection. **(D)** Heavy chain VH family usage in LCMV-specific B cell BCR repertoires from LCMV Armstrong and LCMV clone 13 infected mice at day 45 post-infection. Each point represents one mouse. Data representative of two independent experiments. **(E)** Frequency of mutation in the heavy chain VH genes in LCMV Armstrong and LCMV clone 13 infected mice at d45 post-infection. Data represent VH germline mutation frequency of one representative sequence from each clonotype. Data are pooled from IgH BCR repertoires of 5 individual mice per infection. ***p<0.001, Mann-Whitney U-test of medians. **(F)** Selection pressure (Σ) in IgH CDR and FWR measured by BASELINE algorithm. Data are pooled from IgH BCR repertoires of 5 individual mice per infection. ***p<0.001, Student’s t-test.

Given that accumulation of mutations in immunoglobulin genes is critical for driving affinity maturation (Mesin et al., 2016; Muramatsu et al., 2002; Odegard and Schatz, 2006), we measured heavy chain B cell receptor (BCR) repertoire usage and mutation levels by sequencing BCR genes from LCMV-specific B cells at day 45 p.i. Extensive clonal expansion was observed during both acutely resolved and chronic infection with over 50% of each of the BCR repertoires sequenced represented by the top 20 most abundant clonal families (**Figure S2B**). VH gene family usage was similar between LCMV-specific B cells in acutely resolved and chronic infection and, in both cases, favored the use of the VH1 gene family (60% of all clones), in particular VH1-82 (**Figure 2D** and **S2C**). Despite the similarity in VH gene usage, the accumulation of mutations during acutely resolved and chronic infection was different. LCMV-specific B cells in acutely resolved infection were approximately 2.7% mutated from their germline VH gene sequence, whereas LCMV-specific B cells in chronic infection were only 2% mutated from germline (**Figure 2E** and **S2D**). In chronic infection, decreased BCR mutations occurred preferentially in the complementarity determining regions (CDR) 1 and 2, but not in the framework regions (**Figure S2E**). Additionally, mutations resulting in amino acid replacements were also decreased in the CDR regions during chronic infection (**Figure S2F**). We next applied the BASELINe algorithm (Yaari et al., 2012) to quantify selection pressure in the CDR and framework regions. BCR sequences from LCMV-specific B cells in chronic infection had less positive selection in the CDRs than in acutely resolved infection (**Figure 2F**). Collectively, these data suggested that chronic infection impaired antibody affinity maturation by decreasing the accumulation of affinity-enhancing mutations during somatic hypermutation in the GC.

### Molecular profiles of LCMV-specific GC, PC and MBC are distinct during chronic infection

Chronic infection promotes unique transcriptional programs and differentiation states in CD8+ and CD4+ T cells (Crawford et al., 2014; Doering et al., 2012) but it is unclear whether chronic infection impacts transcriptional programs of antigen-specific B cells and, if so, how. To interrogate this question, we sorted LCMV-specific GC B cells, PC, and MBC at days 7, 15, and 45 p.i. during acutely resolved and chronic infection (**Figure S3A**) and performed bulk RNA-seq. In general, each LCMV-specific B cell population (GC B cells, PC, and MBC) clustered by day p.i. in principal component space (**Figure 3A**). For all three cell types, transcriptional profiles were most similar at day 7 p.i. For PC and MBC, transcriptional programs diverged over time between acutely resolved and chronic infection, with the greatest differences apparent at days 15 or 45 p.i. (**Figure S3B-D**). However, the largest transcriptional divergence between acutely resolved and chronic infection occurred in GC B cells at day 15 p.i. (**Figure 3B**). Indeed, differential gene expression analysis identified 346 genes differentially expressed between LCMV-specific GC B cells at day 15 p.i. Of these 346 differentially expressed genes at day 15 p.i., 311 were upregulated during chronic infection (**Figure 3B and 3C**). In LCMV-specific GC B cells at day 15 p.i., acutely resolved infection enriched for genes related to B cell homeostasis (*Bcl6*, *Lyn*, and *Tnfrsf13c*), cellular response to DNA damage stimulus (*Aicda*, *Foxo1*, *H2afx*, and *Gadd45b*), and histone H3 acetylation (*Brd1* and *Hdac4*) whereas chronic infection enriched for genes related to response to virus (*C1qbp* and *Tlr7*), response to interferon-β and interferon-α (*Irf1*, *Stat1*, *Bst2*, and *Ifi204*), and response to biotic stimulus (*Prdm1*, *Ptpn22*, and *Mif*) (**Figure 3D**). We observed approximately 12-fold increased expression of the PC master regulatory transcription factor *Prdm1* (Turner et al., 1994) and 1.6-fold repression of *Bcl6*, the key transcriptional regulator of GC B cell fate (Crotty et al., 2010), during chronic infection, which suggested that chronic infection may promote excessive LCMV-specific GC B cell differentiation towards the PC fate (**Figure 3E**). Together, these data suggested that LCMV-specific GC B cells have divergent transcriptional programs at day 15 p.i., and expression of key transcription factors pointed to potentially altered GC B cell fate decisions.

**Figure 3:**
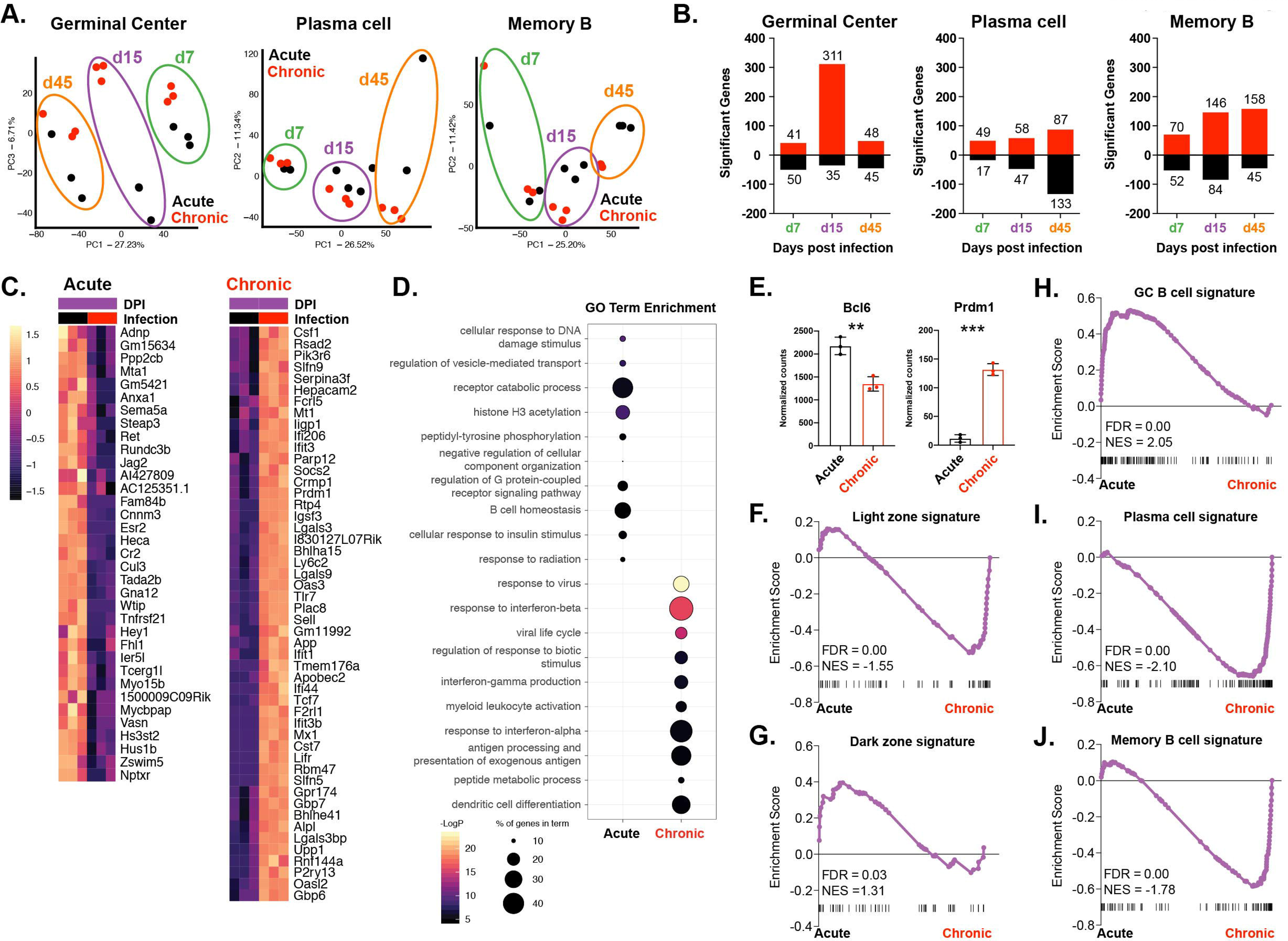
Molecular profiles of LCMV-specific GC, PC and MBC are distinct during chronic infection. **(A)** PCA plot of LCMV-specific GC B cells, PC, and MBC at days 7, 15, and 45 post-infection. **(B)** Number of significant differentially expressed genes (p<0.05 and >2 log fold-change) between LCMV-specific GC B cells, PC, and MBC from LCMV Armstrong (acute) and LCMV clone 13 (chronic) infected mice at the indicated timepoints post-infection. **(C)** Heatmap depicting top 35 and bottom 50 differentially expressed (p<0.05 and >2 log fold-change) genes between LCMV-specific GC B cells from LCMV Armstrong and LCMV clone 13 infected mice at day 15 post-infection. **(D)** Top 10 GO terms enriched in LCMV-specific GC B cells from LCMV Armstrong and LCMV clone 13 infected mice at day 15 post-infection. Color of points represents p-value significance. Size of point represents percent of cluster genes enriched in each respective GO term. **(E)** mRNA normalized counts of *Bcl6* and *Prdm1* in LCMV-specific GC B cells from LCMV Armstrong and LCMV clone 13 at day 15 p.i. **(F)** GSEA plot of light zone gene signature in LCMV-specific GC B cells from LCMV Armstrong and LCMV clone 13 infected mice at day 15 post-infection. **(G)** GSEA plot of dark zone gene signature in LCMV-specific GC B cells from LCMV Armstrong and LCMV clone 13 infected mice at day 15 post-infection. **(H)** GSEA plot of GC B cell gene signature in LCMV-specific GC B cells from LCMV Armstrong and LCMV clone 13 infected mice at day 15 post-infection. **(I)** GSEA plot of PC gene signature in LCMV-specific GC B cells from LCMV Armstrong and LCMV clone 13 infected mice at day 15 post-infection. **(J)** GSEA plot of MB gene signature in LCMV-specific GC B cells from LCMV Armstrong and LCMV clone 13 infected mice at day 15 post-infection.

We focused our analysis on day 15 p.i., which had the greatest transcriptional divergence between infections. The GC is comprised of two anatomical zones where GC B cells undergo selection (light zone; LZ) and accumulate mutations through proliferation (dark zone; DZ) (Mesin et al., 2016). Given the data above on altered accumulation of affinity-enhancing mutations in the BCR during chronic infection, we hypothesized that chronic infection may disrupt the LZ to DZ balance for virus-specific B cells. Using Gene Set Enrichment Analysis (GSEA) (Subramanian et al., 2005), we compared LCMV-specific GC B cells from acutely resolved and chronic infection to known antigen-specific GC B cell LZ and DZ gene signatures (Victora et al., 2010). In chronic infection, LCMV-specific GC B cells were enriched for the LZ gene signature whereas LCMV-specific GC B cells in acutely resolved infection were enriched for the DZ signature (**Figure 3F** and **3G**), suggesting an altered GC distribution of virus-specific B cells in chronic infection. Selection into the DZ is driven by Myc and the mTORC1 complex following positive selection in the LZ (Calado et al., 2012; Dominguez-Sola et al., 2012; Ersching et al., 2017; Finkin et al., 2019; Luo et al., 2018). The mTORC1 signaling gene signature was enriched in chronic infection; however, the Myc target gene signature was unchanged between acutely resolved and chronic infection (**Figure S3E**). Given the critical role for mTORC1 signaling in PC differentiation (Benhamron et al., 2015; Jones et al., 2016) as well as the increased expression of *Prdm1* and decreased expression of *Bcl6* noted in **Fig 3E** during chronic infection, we next compared LCMV-specific GC B cells from acutely resolved and chronic infection to known antigen-specific GC B cell, PC, and MBC gene signatures using GSEA (Bhattacharya et al., 2007; Shi et al., 2015). In agreement with the flow cytometric data in **Fig 1D** and transcription factor analysis in **Fig 3E**, LCMV-specific GC B cells from acute infection enriched for the GC B cell signature but not the PC or MBC signatures (**Figures 3H, 3I**, and **3J**). In contrast, LCMV-specific GC B cells in chronic infection enriched for the PC and MBC cell signatures rather than the GC B cell signature. Collectively, these data indicate that chronic infection promotes skewing toward the PC and MBC cell transcriptional programs within GC B cells, which may indicate early terminal differentiation within the GC in chronic infection.

### Single-cell RNA sequencing of LCMV-specific GC B cells reveals germinal center organization

Our bulk RNA sequencing data suggested divergent fate decisions for LCMV-specific GC B cells responding to acutely resolved or chronic infection including an enrichment in signatures of PC and MBC in the GC population in chronic infection. To examine the GC reaction at increased resolution and dissect the specific steps impacted by chronic infection, we sorted LCMV-specific GC B cells at day 15 p.i. from acutely resolved or chronic infection and performed droplet-based single-cell RNA sequencing (scRNAseq). In total, we analyzed 3270 and 3374 LCMV-specific GC B cells from GCs of acutely resolved and chronic infection, respectively. Unsupervised clustering without consideration of infection type identified seven populations of LCMV-specific GC B cells that clustered according to unique transcriptional and functional profiles (**Figure 4A** and **S4A-B**). We investigated whether the clusters identified could be assigned to the LZ or the DZ. Single-cell enrichment scores were calculated for each cell using previously published genesets for the LZ and DZ GC B cell programs (**Figure 4B** and **S4C**) (Bhattacharya et al., 2007; Shi et al., 2015). Using this approach, we identified enrichment of the LZ gene program in clusters 2, 4, 5, and 6 and enrichment of the DZ gene program in clusters 0, 1, and 3. We next sought to identify the signatures of early PC and MBC, the two cellular outputs of the GC reaction. Calculation of single-cell enrichment scores for previously published PC (Shi et al., 2015) and MBC (Bhattacharya et al., 2007) genesets identified enrichment of the MBC gene program in cluster 5 and the PC gene program in cluster 6 (**Figure 4C** and **S4D**). Enrichment of the PC and MBC transcriptional programs occurred within clusters that also enriched for the LZ program, supporting previously published observations (Ise et al., 2018; Kräutler et al., 2017; Laidlaw et al., 2017; Suan et al., 2017) that the decision to undergo terminal differentiation and exit the GC occurs within the LZ.

**Figure 4:**
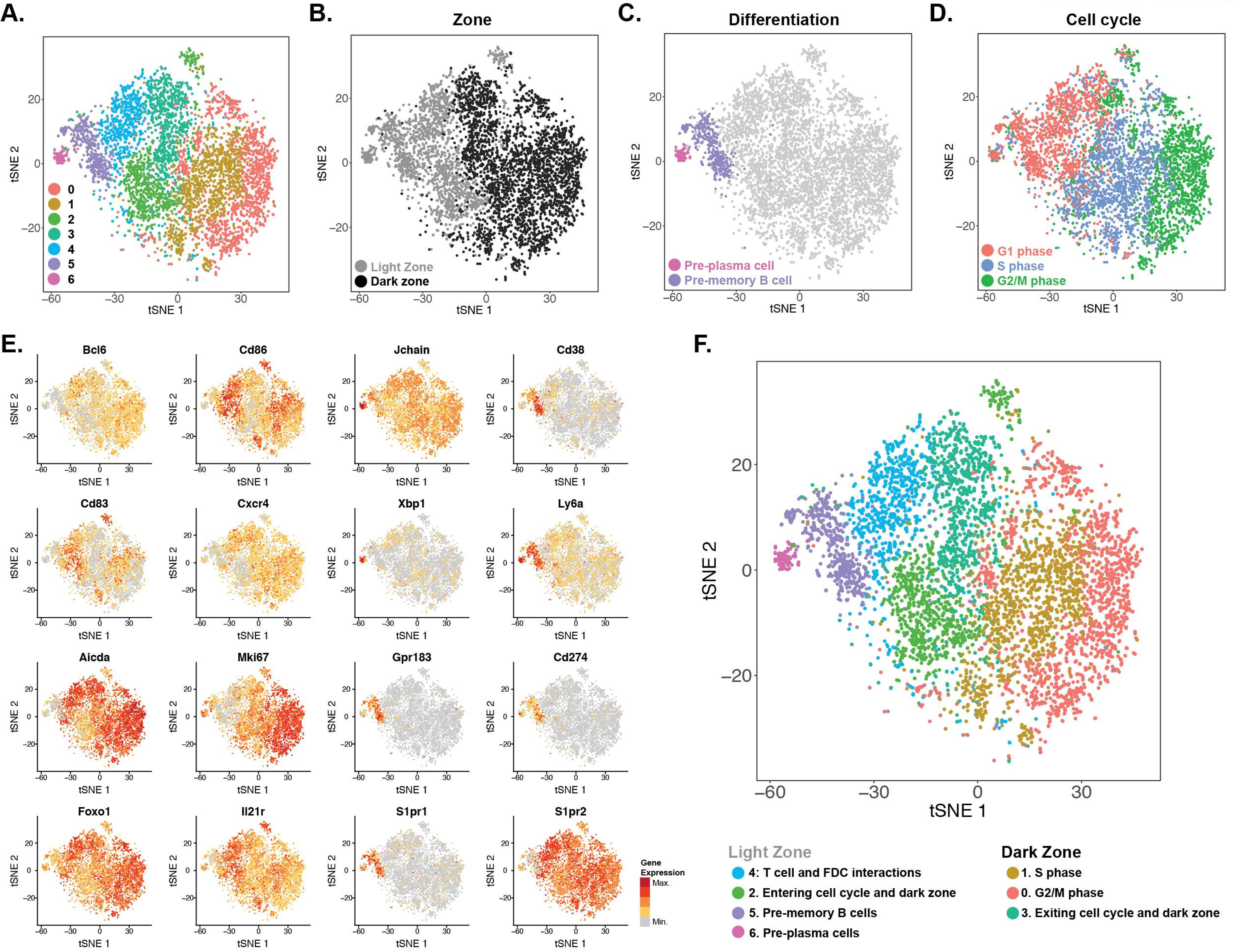
Single-cell RNA sequencing of LCMV-specific GC B cells reveals germinal center organization. **(A)** tSNE plot of unbiased clustering of LCMV-specific GC B cells at day 15 post-infection. **(B)** tSNE plot depicting single-cell light zone or dark zone geneset enrichment. **(C)** tSNE plot depicting single-cell MB or PC geneset enrichment. **(D)** tSNE plot depicting cell cycle stage enrichment scores for each cell. **(E)** Plots showing expression of selected genes important for GC B cell biology. **(F)** tSNE plot depicting cluster identification and cluster annotation of LCMV-specific GC B cells determined using unbiased clustering.

Given the importance of cell proliferation in the GC reaction (Gitlin et al., 2014, 2015; Mcheyzer-Williams et al., 2015), we next investigated the role of the cell cycle in driving clustering of LCMV-specific GC B cells. Consistent with cell cycle biology as a major driver of the GC reaction, we observed restriction of active proliferation and enrichment for the S phase and G2/M phase of the cell cycle in clusters belonging to the DZ (clusters 0, 1, and 3) (**Figure 4D**). In fact, the majority of heterogeneity observed within the DZ clusters was driven by differences in cell cycle stage, with cluster 0 enriching for cells in the G2/M phase, cluster 1 enriching for cells in the S phase, and cluster 3 containing a mixture of cells from all three phases of cell cycle (**Figure S4E**). In contrast, the majority of cells in clusters 4, 5, and 6 were in G1, indicating that these cells had exited from active division. These data are consistent with previous studies (Dominguez-Sola et al., 2012; Finkin et al., 2019; Gitlin et al., 2015) that identified regulation of cell cycle entry as critical for the GC reaction. In line with the calculated single-cell enrichment scores in **Figure 4B-D**, we observed cluster- and zone-restricted expression of individual genes central to the LZ (*Cd83, Cd86*) or the DZ programs (*Mki67, Cxcr4*) (**Figure 4E**) (Victora et al., 2010, 2012). Notably, despite being in cell cycle, clusters 2 and 3 enriched for the LZ and DZ signatures, respectively. These data, combined with the cell cycle analysis in **Figure 4D**, suggested that cells in cluster 2 were leaving the LZ, initiating cell cycle, and entering the DZ whereas cells in cluster 3 were leaving cell cycle and exiting the DZ to return to the LZ. Cells in cluster 4 enriched for genes related to antigen processing and presentation via MHCII (*Cd74* and *H2-DMa*) and B cell survival and signaling genes (*Cd79b* and *Tnfrsf13c*) (**Figure S4B**), suggesting that cells in this cluster are interacting with Tfh and/or FDCs (Mesin et al., 2016). In addition to enriching for antigen processing and presentation via MHCII, cluster 5 enriched for genes involved in T cell trafficking, further suggesting that cells in this cluster are exiting the GC as MBC (**Figure S4B**). Cluster 6 was defined by genes central to PC function, including high expression of the PC immunoglobulin secretory factor *Jchain* and genes related to protein folding, protein localization to the ER, and response to ER stress (**Figure S4B**). Other key GC B cell genes such as *Gpr183, Foxo1, Il21r, S1pr1,* and *S1pr2* were expressed in the expected LZ and/or DZ clusters (**Figure 4E**). Thus, single-cell analysis of LCMV-specific GC B cells provided increased resolution into the transcriptional coordination of GC B cell responses and identified cells in the LZ and DZ, the transition between these two phases, and GC B cells undergoing differentiation towards PC or MBC.

### Chronic infection promotes terminal B cell differentiation and GC exit

Although the scRNAseq analysis above provided increased resolution into the GC reaction, this clustering analysis was generated with GC B cells from both acutely resolved and chronic LCMV. Therefore, we next asked how chronic infection alters the distribution of GC B cells and specifically how fate decisions are affected. To this end, we first determined the frequency of LCMV-specific GC B cells from acutely resolved or chronic infection within each cluster. Both acutely resolved and chronic infection had similar frequencies of LCMV-specific GC B cells in the DZ clusters (**Figure 5A**). However, chronic infection resulted in increased frequency of LCMV-specific GC B cells in the pre-PC (cluster 6) and pre-MBC (cluster 5) clusters by 6.6-fold and 1.8-fold, respectively. Conversely, chronic infection diminished the frequency of cells in transition from LZ to DZ (cluster 2) by 0.7-fold. Thus, chronic infection promoted accumulation of virus-specific GC B cells in the pre-PC and pre-MBC fates and depletion of cells re-entering the cell cycle and the DZ.

**Figure 5:**
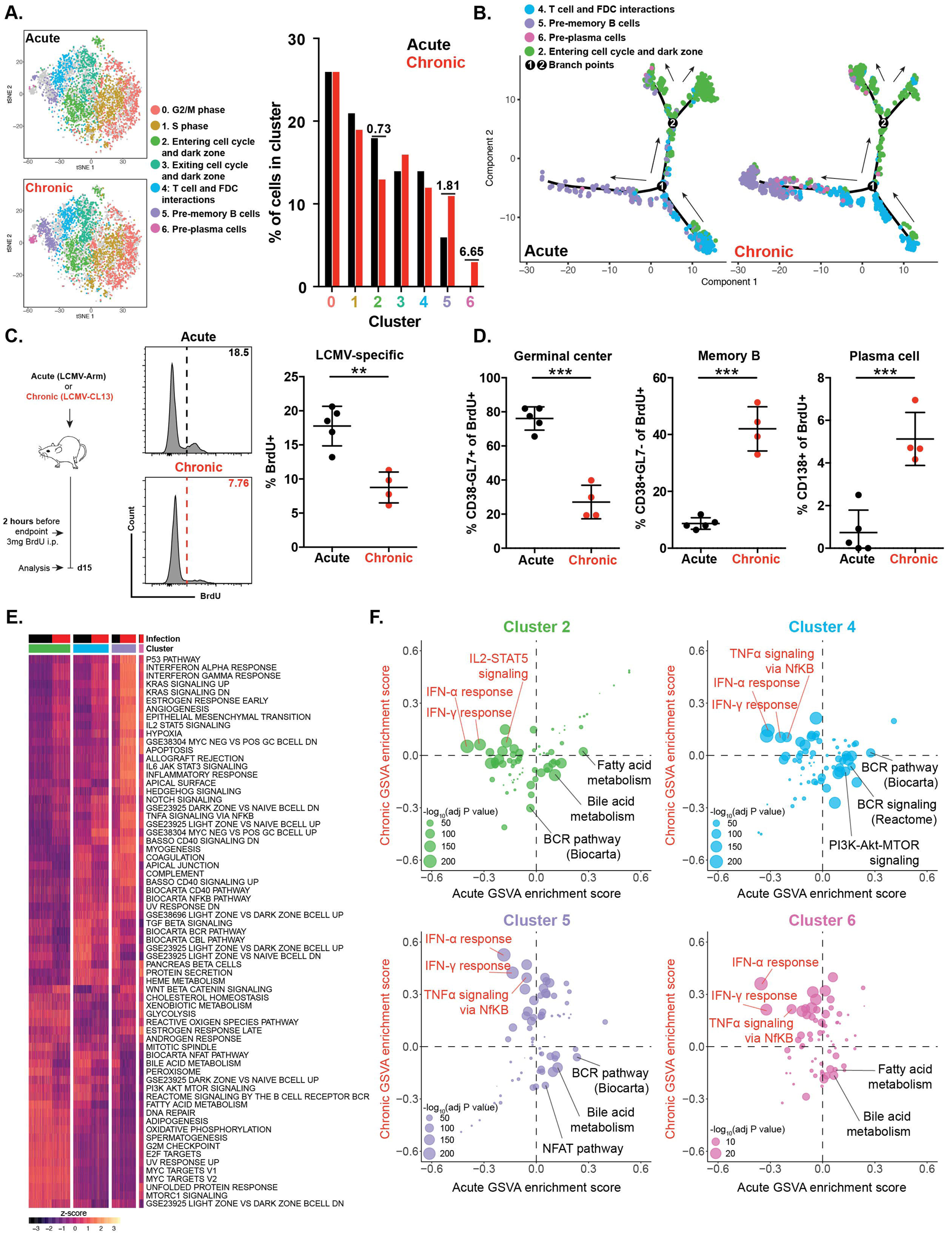
Chronic infection promotes terminal B cell differentiation and GC exit. **(A)** tSNE and bar plots depicting cluster occupancy of LCMV-specific GC B cells from LCMV Armstrong (Acute) and LCMV clone 13 (Chronic) infected mice at day 15 post-infection. **(B)** Pseudotime analysis of light zone GC B cell populations identified using scRNAseq. Pseudotime constructed using all light zone GC B cell populations and visualized separated by infection. Pseudotime is rooted in cluster 4: T cell and FDC interactions (see Figure S5A). **(C)** Design of short-term BrdU labeling experiments. Mice were infected with either acute or chronic LCMV and treated with 3mg/mouse of BrdU 2 hours prior to analysis at day 15 post-infection. Representative histograms of BrdU labeling of LCMV-specific GC B cells from acute or chronic LCMV infection (left) and summary data of BrdU labeling in LCMV-specific B cells (right). **(D)** Frequency of BrdU+ LCMV-specific GC B cells, memory B cells, and plasma cells at day 15 post-infection according to experimental design in (C). **(E)** Heatmap of single-cell GSVA geneset enrichment scores for all LZ clusters from LCMV-specific GC B cell scRNAseq dataset. **(F)** Bubble plots of acute vs chronic single-cell GSVA enrichments scores for each LZ cluster. Point size reflects p value. Selected differentially expressed genesets are labeled. Data points in C and D show mean ± SEM from 4-5 mice from 1 of 2 representative experiments. **p<0.01, ***p<0.001, Student’s t-test.

It remained unclear where chronic infection caused diversion in the stepwise, cyclic scheme of GC B cell differentiation. A typical progression through the GC involves rapid proliferation and somatic hypermutation in the DZ, exit from cell cycle and testing of BCR, and then key decision points in the LZ to initiate either DZ re-entry or begin a PC or MBC differentiation path (Mesin et al., 2016; Shlomchik et al., 2019; Stewart et al., 2018). To understand how chronic infection altered these biological decision points in the GC, we next applied pseudotime analysis to the single-cell data to infer differentiation trajectories and cluster relationships (Qiu et al., 2017b, 2017a). Pseudotime analysis of combined infection types revealed two major branch points (**Figures 5B** and **S5A**). The first branch point (branch point 1) was between GC B cells interacting with FDC and Tfh (cluster 4), GC B cells returning to the DZ (cluster 2), and GC B cells undergoing differentiation into the pre-MBC and pre-PC fates (clusters 5 and 6, respectively). LCMV-specific GC B cells in chronic infection favored the branch toward the pre-MBC and pre-PC fates whereas cells in acutely resolved infection favored the DZ re-entry branch (**Figure 5B**). This pseudotime analysis was consistent with the strong shift in cluster frequency seen in **Figure 5A**, in which LCMV-specific GC B cells from chronic infection were highly biased toward the pre-PC and pre-MBC cell fates. The second branch point (branch point 2) that occurred in the DZ re-entry cluster also separated cells by infection (**Figure 5B**). These data suggested that chronic infection not only drives terminal differentiation of LZ GC B cells during interactions with Tfh and FDC, but that chronic infection also alters LZ GC B cell re-entry to the DZ.

To test the pseudotime prediction that GC B cells in chronic infection would skew differentiation toward the PC and MBC fates more than the DZ re-entry observed in acutely resolved infection, we treated mice infected with LCMV Arm or clone 13 with 5-bromo-2’-deoxyuridine (BrdU) 2 hours prior to analysis (**Figure 5C**). Because BrdU is only incorporated into actively dividing cells, the short BrdU labeling prior to analysis would provide a defined window into the cellular output of the GC reaction at day 15 p.i. The frequency of cells incorporating BrdU was 2-fold lower in LCMV-specific B cells in chronic infection (**Figure 5C**), indicating less proliferation overall. Of the LCMV-specific B cells that did proliferate during the 2hr labeling window (BrdU+), there were lower GC B cell frequencies and higher MBC and PC frequencies in chronic infection compared to acutely resolved infection (**Figure 5D**). Although some BrdU incorporation in MBC and PC could have occurred outside the GC, these data were nonetheless consistent with the scRNAseq pseudotime analysis and further supported the notion that chronic infection skewed LCMV-specific GC B cells towards terminal differentiation and GC exit rather than DZ re-entry.

We next sought to investigate the signals that caused GC B cells to skew toward terminal differentiation during chronic infection using gene set variation analysis (GSVA) to analyze scRNAseq data at the pathway level (Hänzelmann et al., 2013). GSVA identified many pathways that were differentially enriched in LCMV-specific GC B cells between acutely resolved and chronic infection. We projected key GSVA signatures in pseudotime space (**Figure S5B-G**) and directly compared GSVA enrichment scores in acutely resolved versus chronic infection (**Figure 5E-F**).

Signals from T follicular helper cells (Tfh) are critical during the cluster 4 testing phase and chronic LCMV infection leads to an abundance of Tfh (Crawford et al., 2014; Fahey et al., 2011; Vella et al., 2017). Indeed, several Tfh-derived signals were implicated in the scRNA-seq and bulk RNA-seq above. To understand the role of Tfh signaling within the context of GC B cell fate decisions, we analyzed signals derived from Tfh cell help, given that Tfh cell help to GC B cells occurs via pathways including MHCII:TCR (Crotty, 2015; Schwickert et al., 2011) and CD40-CD40L interaction that drive GC B cell survival and entry into the DZ (Ise et al., 2018; Kawabe et al., 1994; Watanabe et al., 2017). Both acute and chronic infection upregulated expression of genes related to the response to CD40 signaling within LZ GC B cells of cluster 4 and along both the branch leading to PC and MBC differentiation as well as the DZ re-entry branch (**Figure S5B**). This result was consistent with abundant Tfh during both LCMV Arm and clone 13 infections (Crawford et al., 2014; Fahey et al., 2011; Vella et al., 2017) and that CD40 agonism can drive both GC B cell DZ re-entry and terminal PC differentiation *in vitro* (Erickson et al., 2002; Foy et al., 1996). However, although the enrichment was similar between infections, the pattern of this CD40 help signal was distinct. Enrichment of CD40 signaling genes in acutely resolved infection increased early in the pseudotime trajectory. In chronic infection, this CD40 signature was instead predominantly found in the latter parts of the pseudotime trajectory, where the pre-MBC and pre-PC features were present. Thus, despite the abundance of Tfh cells in chronic infection (Crawford et al., 2014; Fahey et al., 2011; Vella et al., 2017), these data suggested that Tfh help may not be available during the same steps in GC B cell differentiation. Whether these differences are due to B cell intrinsic features, alterations in Tfh homing and migration, GC architectural differences, or help-signal delivery will require future examination.

Following interaction with Tfh cells and positive selection in the GC, GC B cells upregulate metabolic pathways, Myc targets, and MTOR signaling in anticipation of re-entering cell cycle and the DZ (Dominguez-Sola et al., 2012; Ersching et al., 2017; Finkin et al., 2019; Luo et al., 2018). Because LCMV-specific GC B cells in acutely resolved versus chronic infection appeared to differ at the DZ re-entry step (**Figure 5B**) and because CD40 signals were diminished in the T cell and FDC interaction cluster during chronic infection, we hypothesized that induction of MTOR or Myc target genes would be reduced in chronic infection. However, in contrast to our hypothesis, Myc target genes and MTORC1 signaling were enriched in LZ GC B cells returning to the DZ during both acutely resolved and chronic infection, and neither pathway was differently associated with the observed branching between the two infections (**Figure 5B, S5D,** and **S5E**), though the PI3K-Akt-MTOR hallmark geneset was biased towards stronger enrichment in acutely resolved infection (**Figure S5C**). These observations suggested that, once an effective BCR and help signal was received, engagement of the downstream metabolic transcriptional program proceeded similarly in acutely resolved versus chronic infection.

In contrast to the relatively similar metabolic signatures, we observed positive enrichment of type I interferon signaling, interferon-γ signaling, and TNF signaling in all LZ GC B cell clusters during chronic infection but negative enrichment of these pathways in the same clusters during acutely resolved infection (**Figure 5F, S5F**, and **S5G**), consistent with our bulk RNAseq data (**Figure 3D**). Enrichment of type I interferon, interferon-γ signaling, and TNF signaling was increased in the pre-PC (cluster 6) and pre-MBC (cluster 5) clusters, suggesting that inflammatory signaling may play a role in skewing GC B cell differentiation away from the DZ and towards GC exit during chronic infection. Moreover, even in the second pseudotime branch point at DZ re-entry, inflammatory signals showed a bias in the branch that had a lower signal for PI3K-Akt-MTOR signatures during chronic infection (**Figure S5C**), suggesting an anti-correlation between inflammatory signaling and BCR or metabolic signaling. Indeed, BCR signaling pathways were enriched in acutely resolved infection and strongly anti-correlated with inflammatory signaling (**Figure 5F**). In particular, BCR signaling and PI3K-Akt-MTOR signaling was enriched during acutely resolved infection in cluster 4, where GC B cells were undergoing testing and positive selection, and within cluster 5, the pre-MBC cluster (**Figure S5C**). Conversely, BCR and PI3K-Akt-MTOR signaling were negatively enriched in these same clusters during chronic infection. Chronic infection was also associated with negative enrichment of BCR pathway genes in GC B cells returning to the DZ (cluster 2), where positive enrichment was observed in acutely resolved infection. Together, these data suggested that GC fate decisions in chronic infection were associated with both persistent inflammation and altered sensing of antigen. In particular, the effects of inflammation and BCR signaling were most marked during a key step in the transition from LZ to DZ (i.e. cluster 4). In this testing cluster, BCR signaling attenuation and inflammatory signaling may differentially instruct cells as they diverge between re-entering the DZ or initiating the GC exit program towards the PC or MBC fate.

### Chronic inflammation and altered antigen sensing skew GC B cell responses during chronic viral infection

GSVA identified inflammatory and BCR signaling as two of the most divergent pathways between acutely resolved and chronic infection. Although differences in inflammatory signaling were expected, attenuation of BCR in chronic infection was surprising given the abundance of antigen (**Figure 5F**). In pseudotime projections of both acutely resolved and chronic infection, BCR signaling signatures were enriched in cells re-entering the DZ but not in cells exiting the GC reaction (**Figure 6A**), supporting the known role for BCR signaling in efficient DZ re-entry (Luo et al., 2018). Although the LCMV-specific GC B cells that did re-enter the DZ during chronic infection displayed modest enrichment for BCR signaling, the enrichment of this signature was lower in magnitude during chronic infection and temporally delayed in pseudotime space compared to acutely resolved infection. Expression of cell surface inhibitory receptors or intracellular signaling attenuators did not provide an obvious explanation for attenuated BCR signaling during chronic infection (**Figure S6A** and **S6B**). Instead, we observed downregulation of several BCR pathway genes in LCMV-specific GC B cells within cluster 4 during chronic infection, including *Cd79b* and *Blnk* (**Figure 6B**). Because the ability to elicit Tfh help and receive CD40 signals were maintained, these results suggested that the BCR signaling was impaired while its endocytic function remained intact.

**Figure 6:**
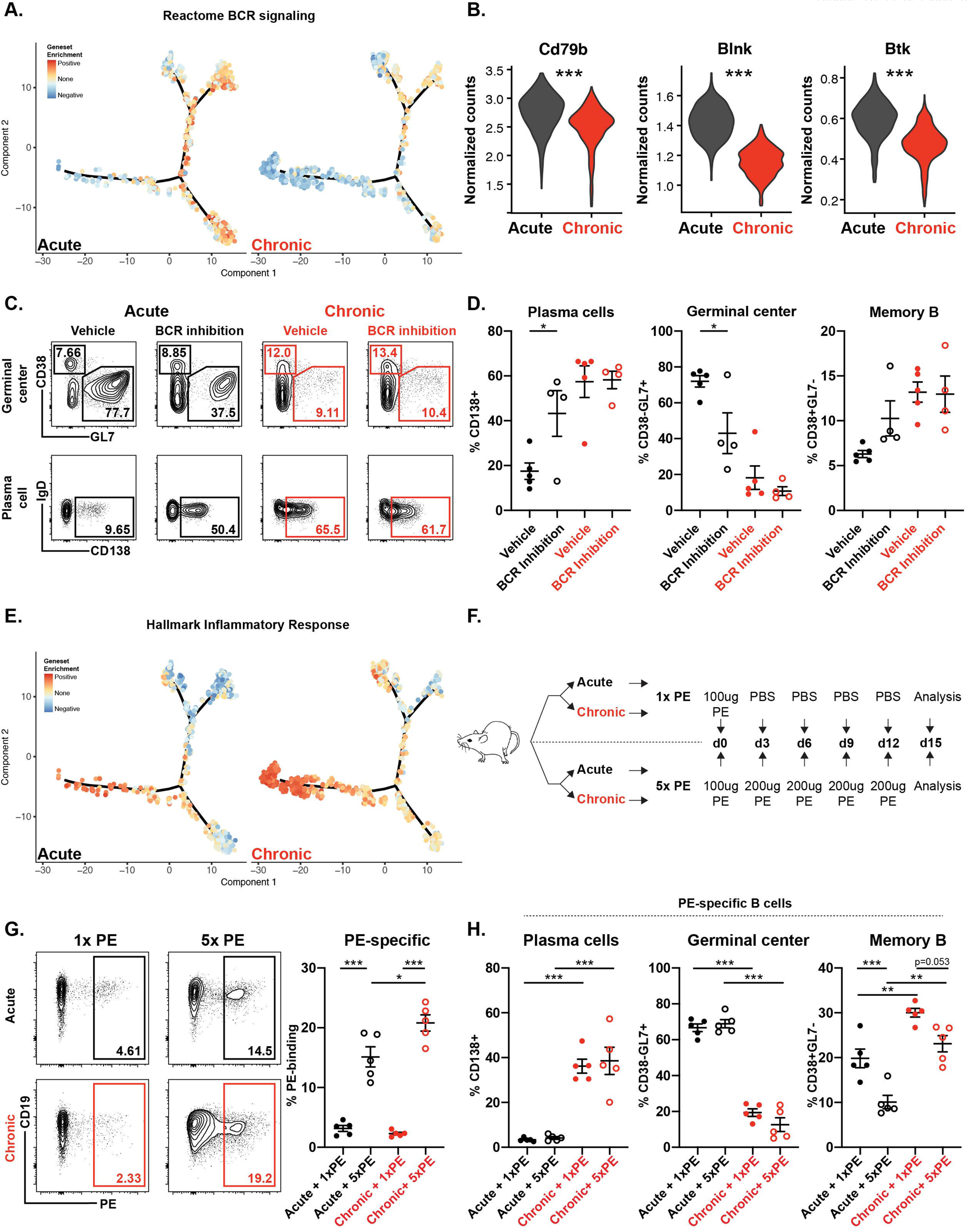
Chronic inflammation and altered antigen sensing skew GC B cell responses during chronic viral infection. **(A)** Single-cell GSVA enrichment scores for Reactome BCR signaling geneset plotted as a function of pseudotime. **(B)** Expression of *Cd79b*, *Blnk*, and *Btk* in LCMV-specific GC B cells within cluster 4 (T and FDC interactions) during LCMV Armstrong (acute) and LCMV clone 13 (chronic) infection at day 15 post-infection. **(C)** Representative FACS plots of GC B cells, memory B cells, and plasma cells from acute and chronic LCMV-infected mice at day 15 post-infection. Mice received either vehicle or ibrutinib twice daily beginning at day 12 post-infection to inhibit BCR signaling. **(D)** Frequency of LCMV-specific plasma cells, GC B cells, and memory B cells at day 15 post-infection from mice receiving BCR inhibition beginning at day 12 as described in C. **(E)** Single-cell GSVA enrichment scores for Hallmark Inflammatory geneset plotted as a function of pseudotime. **(F)** Experimental design of single-dose (1xPE) and chronic dosing (5xPE) experiments. Briefly, mice were immunized with 100ug PE at the time of infection with either LCMV Armstrong or LCMV clone 13. Mice received either 200ul PBS or 200ug PE i.p. at days 3, 6, 9, and 12 post-infection prior to analysis at day 15 post-infection. **(G)** Representative FACS plots of PE-specific B cells in mice as treated in F (left). Frequency of PE-specific B cells in mice as treated in F (right). **(H)** Frequency of PE-specific plasma cells, GC B cells, and memory B cells in mice treated as in F at day 15 p.i. Each plot in D, G, and H depicts mean ± SEM with 4-5 mice per group. *p<0.05, **p<0.01, ***p<0.001, One-way ANOVA followed by Tukey’s multiple comparison test.

Given our observation that BCR signaling was enriched in GC B cells undergoing DZ re-entry during acutely resolved infection (**Figure 6A** and **6B**), we hypothesized that inhibiting BCR signaling during acutely resolved infection would lead to GC B cell exit and differentiation. In contrast, because the BCR signaling transcriptional network was diminished in chronic infection, we predicted that BCR inhibition would have less of an impact in chronic infection. To test these predictions, we administered the BCR signaling (i.e. BTK)-inhibitor ibrutinib to mice infected with acutely resolved or chronic LCMV starting at day 12 p.i. as previously described (Mueller et al., 2015; Yang et al., 2016). Ibrutinib treatment during acutely resolved infection decreased LCMV-specific GC B cell responses and increased PC frequencies, consistent with the known role of BCR signaling in GC re-entry (**Figure 6C** and **6D**). In contrast, inhibition of BCR signaling during chronic infection had no impact on LCMV-specific B cell differentiation. These results were consistent with our scRNAseq analysis in **Figures 5F** and **6A** and suggested that skewing of GC B cell differentiation toward PC and MBC differentiation and away from DZ re-entry during chronic infection was associated with attenuated BCR signaling. Moreover, such an impact on the GC response could be partially recapitulated by molecular BCR inhibition with ibrutinib.

Chronic infections can not only impact antibody responses to the persisting infection itself, but also bystander antigens such as vaccines (McKittrick et al., 2013; Shive et al., 2018; Wiedmann et al., 2000). This observation suggests a role for the chronic inflammatory environment (Stelekati and Wherry, 2012), in addition to persisting antigen, in altering B cell responses to chronic infection. Thus, we next sought to determine the role of persistent inflammation in altering GC B cell differentiation. Genes related to inflammatory responses or downstream of interferon-α and interferon-γ were strongly enriched in LCMV-specific GC B cells undergoing differentiation into PC and MBC (**Figure 6E, S5F,** and **S5G**). Moreover, inflammatory signaling gene transcription was also enriched even in the pseudotime branch of LZ GC B cells returning to the DZ that was biased to chronic infection. Thus, persistent inflammatory signaling during chronic infection was strongly associated with the skewed differentiation of LCMV-specific GC B cell towards the PC and MBC and early GC exit.

To test whether the inflammatory environment of chronic infection impacted the GC B cell response, we next developed a more pathway-agnostic experimental approach. We infected mice with acutely resolved or chronic LCMV and then injected these mice with one dose of recombinant phycoerythrin (1x PE; **Figure 6F**) (Pape et al., 2011). We then assessed the impact of the inflammatory environment of chronic infection on the PE-specific B cell response. We hypothesized that persistent inflammation during chronic infection would skew PE-specific GC B cells towards PC and MBC differentiation. Although the magnitude of the overall PE-specific B cell response was similar between acutely resolved and chronic LCMV-infected mice (**Figure 6G**), the differentiation of PE-specific B cell responses in mice with chronic infection was altered (**Figure 6H**). Indeed, the frequencies of PE-specific PC were increased during chronic infection compared to acutely resolved infection, indicating a role for environment and/or inflammation in the skewing of B cell responses during chronic infection towards PC differentiation. These data are consistent with the GC B cell inflammatory signatures and the bias of GC output during chronic infection towards the PC fate.

The effects of inflammatory signals during chronic infection impacted LCMV-specific and bystander specificities. It was unclear, however, whether the observed BCR signaling alterations were related to excessive antigen exposure or whether other factors such as inflammation or environment were the main drivers of altered humoral immunity in chronic infection. To test the potential consequences of attenuated BCR signaling, we designed a variation of the PE immunization experiment that used a repeated immunization scheme to expose B cells to antigen “chronically” for 2 weeks (**Figure 6F**). Repeated administration with PE increased the magnitude of the PE-specific B cell response in both acutely resolved and chronic infection (**Figure 6G**). Repeated immunization with PE did not, however, impact PE-specific PC frequencies in either infection model, suggesting that increased antigen dose and duration were not able to impact the PC fate during LCMV (**Figure 6H**). Similarly, repeated immunization also did not alter the frequency of PE-specific GC B cells in either infection model, and the attenuated GC B cell response observed in chronic infection was not improved by repeated immunization (**Figure 6H**). While PC and GC B cell responses were not altered by “chronic” antigen exposure, the MBC compartment in both acutely resolved and chronic infection was decreased in magnitude (**Figure 6H**). These experiments suggest that increasing antigen alone in acutely resolving infection is not sufficient to drive the skewed B cell fate observed during chronic infection and that the inflammatory environment in chronic infection plays a dominant role in driving an aberrant GC B cell fate.

## DISCUSSION

Humoral immune responses during chronic infections are often dysregulated and characterized by hypergammaglobulinemia, decreased affinity maturation, and delayed development of neutralizing antibodies (Battegay et al., 1993; Bjøro et al., 1994; Cashman et al., 2014; Eschli et al., 2007; Gray et al., 2007; Hunziker et al., 2003; Lane et al., 1983; Logvinoff et al., 2004; Milito et al., 2004; Moir and Fauci, 2017; Oldstone and Dixon, 1969; Racanelli et al., 2006). Prior studies have suggested that poor antibody quality is in part due to deletion of B cells prior to establishment of the GC response (Fallet et al., 2016; Moseman et al., 2016; Sammicheli et al., 2016). Our data extend these findings to show that some LCMV-specific B cells do establish a GC during chronic infection; however, the dynamics of the GC response in chronic infection are dysregulated and this dysregulation is associated with poor antibody quality and hypergammaglobulinemia. Detailed interrogation of LCMV-specific GC B cells identified the molecular and cellular events associated with this dysregulation, providing insights into a previously largely unexplored aspect of altered humoral immunity during chronic infection. In this study, we found that chronic infection skews virus-specific GC B cell responses towards terminal PC and MBC differentiation and away from continued participation in the GC response. Moreover, this skewing is associated with early GC B cell exit during chronic infection due to an altered GC B cell decision point in the LZ. This skewing is associated with decreased affinity maturation, increased inflammatory signaling, and attenuated BCR signaling. In addition, we identified a role for the environment of chronic infection in perturbing the GC response; these alterations also impacted B cell responses to antigen that are unrelated to the persisting virus, implicating these GC effects in broader consequences of bystander chronic infections. These data provide not only a detailed single-cell molecular and transcriptional map of GC B cell responses following acute infection, but also insights into molecular programs of GC B cells that are altered by persisting infection.

To define the point of altered GC B cell fate decisions during chronic infection, we interrogated LCMV-specific GC B cell responses using cellular approaches as well as bulk and single-cell transcriptional profiling. Bulk RNA-seq identified day 15 p.i. as the most transcriptionally distinct timepoint between virus-specific B cells in acutely resolved versus chronic infection, especially for GC B cells. By focusing scRNAseq at this time point on the GC and employing pseudotime analysis, we were able to resolve the cyclic GC B cell reaction and individual steps in the LZ and DZ. This high temporal resolution revealed fairly similar biology in the DZ in acutely resolved versus chronic infection, but several clear differences in the LZ. Specifically, the decision to undergo PC and MBC differentiation was identified within the LZ for both acutely resolved and chronic infection – a result consistent with other recently published studies (Ise et al., 2018; Kräutler et al., 2017; Laidlaw et al., 2017; Suan et al., 2017). However, this key checkpoint was altered during chronic infection with an increase in GC exit, differentiation to pre-MBC and pre-PC, and diminished DZ re-entry. Pathway-level analysis revealed attenuated BCR signaling and increased inflammatory signaling at this decision point that favored GC exit over DZ re-entry. Dysregulation of humoral immunity during chronic infection has been linked to pre-GC B cell deletion (Fallet et al., 2016; Moseman et al., 2016; Sammicheli et al., 2016) and excess Tfh (Crawford et al., 2014; Fahey et al., 2011; Vella et al., 2017), but the B cell intrinsic events in the GC have not previously been defined. The single-cell nature of these studies now identify not only a role for altered GC biology, but also the specific decision point in the GC that is altered by chronic infection. Moreover, these studies revealed B cell-intrinsic effects of chronic infection, including a potential antagonistic relationship between inflammatory signaling and BCR signaling in GC B cells.

Attenuated BCR signaling during chronic infection in the context of abundant antigen was revealed in these studies. This observation is, at least superficially, consistent with the notion of overstimulation and/or negative regulation of lymphocyte responses during chronic infections (McLane et al., 2019; Vella et al., 2017). Although B cell signaling could be attenuated by many different types of cell surface receptors, GC B cells during chronic LCMV infection did not display obvious upregulation of canonical B cell inhibitory receptors. Changes in BCR signal strength can, however, have implications for GC B cell fate (Ochiai et al., 2013; Phan et al., 2006; Turner and Ke, 2018). Current models of GC B cell differentiation indicate that high strength of signal received in the LZ promotes PC differentiation whereas low strength of signal fosters MBC differentiation or death (Ise and Kurosaki, 2019; Shlomchik et al., 2019). Our data are generally consistent with the notion that high antigen signal strength favors PC differentiation but also reveal several additional mechanistic insights into GC biology during chronic infection. For example, our data reveal BCR signaling attenuation during chronic infection and point to two underlying molecular circuits. First, BCR signaling molecules are downregulated during chronic infection. Together with essentially normal signatures for CD40:CD40L signaling, this observation raises the possibility that, whereas BCR signaling potential is lower, the ability to capture and present antigen to Tfh and receive help in the form of CD40 signals remains intact. These data might link to previous observations about the role of excess Tfh help during chronic infections (Crawford et al., 2014; Fahey et al., 2011; Vella et al., 2017). Second, our data reveal a potential relationship between inflammatory signaling and altered BCR signaling. In contrast to settings where higher antigen stimulation fosters PC differentiation, the high antigen load and excessive PC formation during chronic viral infection was associated with attenuated rather than stronger BCR signaling. Moreover, this enhanced PC production was not disrupted by small molecule inhibition of BCR signaling. Attenuated BCR signaling in the GC exit branch of the pseudotime was also associated with robustly increased signatures of inflammatory signaling. These data suggest that inflammation plays a role in altered GC B cell fate decisions, including driving the major difference in the LZ, where GC B cells are guided to either re-enter the DZ or exit the GC as PC or MBC. Even in the DZ re-entry branch of the pseudotime analysis, signatures of inflammatory signaling and BCR signaling split into two subbranches. High antigen loads and excessive BCR triggering could undoubtedly contribute to signaling desensitization. However, these new data reveal a picture of GC B cell biology during chronic infection where inflammatory, BCR, and help signals from Tfh are integrated to create a distinct equilibrium that favors GC exit over re-entry that likely contributes to both lower quality antibody and hypergammaglobulinemia.

The scRNA-seq (and bulk RNA-seq) revealed several distinct pathways of inflammation, but a role for interferon signaling in shaping the humoral immune response was particularly prominent. Prior studies suggested a major role for interferon before the GC develops but antigen-specific B cells were previously thought to be insensitive to type I interferon after entry into the GC reaction, given their low expression of cognate cytokine receptors (immgen.org). Here, we identified a strong transcriptional imprint of interferon signaling on the GC B cells. This signal enriches particularly in cells at the key decisional node for DZ re-entry versus GC exit. It is unclear, however, whether this transcriptional signature is due to direct IFNAR signaling, IFNγR signaling, or perhaps other signaling mechanisms such as TLR7 (Clingan and Matloubian, 2013; Walsh et al., 2012). It is also unclear whether this transcriptional imprint is due to active signaling within the GC or represents a residual signal from interactions occurring before entry into the GC. Nevertheless, the dysregulated or elevated inflammatory signaling in GC B cells during chronic infection may be one mechanism by which chronic viral infections impact immunity to unrelated vaccines or infections (McKittrick et al., 2013; Shive et al., 2018; Wiedmann et al., 2000). Such bystander effects on vaccine immunity are a major challenge for patients with chronic infections or inflammatory diseases, particularly in the developing world where multiple endemic infections are common (Stelekati and Wherry, 2012). Thus, efforts aimed at re-balancing the appropriate cytokine and inflammatory circuits in the GC may reveal therapeutic opportunities to enhance humoral immunity during chronic infections.

Taken together, our studies identify a key GC B cell differentiation checkpoint that is dysregulated during chronic infection. Further, we find that the chronic inflammatory environment, rather than persistent antigen, is sufficient to drive altered GC B cell differentiation during chronic infection even against unrelated antigens. However, our data also indicate that inflammatory circuits are likely linked to perception of antigen stimulation. Here, we have focused on B cell- and GC-intrinsic features of dysregulated humoral immunity during chronic viral infection. It is likely that other factors including immune complexes (Wieland et al., 2015; Yamada et al., 2015), altered stromal cell biology (Mueller et al., 2007), excess or altered Tfh help (Crawford et al., 2014; Fahey et al., 2011; Greczmiel et al., 2017; Vella et al., 2017), regulation of spatial localization, and structural alterations to lymphoid tissues (Matter et al., 2011; Mueller et al., 2007; Wilson et al., 2013) will have a role. Nevertheless, this study reveals a B cell-intrinsic program of transcriptional skewing in chronic viral infection that results in shunting out of the cyclic GC B cell process and early GC exit with consequences for antibody quality and hypergammaglobulinemia. These findings have implications for vaccination in individuals with pre-existing chronic infections where antibody responses are often ineffective and suggest that modulation of inflammatory pathways may be therapeutically useful to overcome impaired humoral immunity and foster affinity maturation during chronic viral infections.

## ACKNOWLEDGEMENTS

We thank members of the Wherry and Luning Prak laboratories for discussions and critical review of the data and manuscript; members of the Allman laboratory (UPenn) and Flow Cytometry Shared Resource for technical advice regarding cell sorting experiments; the University of Pennsylvania Laboratory Animal Resources and the BRB II/III animal facility, and D. Allman, M. Cancro, C. Hunter, D. Douek, J. Henao-Meija, and B. DeKosky for critical scientific advice. RPS was supported by a NIAID T32 program Training in HIV pathogenesis (T32 AI007632-18). The Human Immunology Core is supported by P30 CA016520 and P30 AI0450080. ETLP and WM are supported by P01 AI106697. LAV was supported by the Pediatric Infectious Diseases Society Fellowship Award, the National Center for Advancing Translational Sciences of the NIH (KL2TR001879), and a Mentored Research Scholar Award from the Penn Center for AIDS Research (CFAR), an NIH-funded program (P30 AI 045008), and a Mentored Clinical Scientist Career Development Award from the National Institute of Allergy and Infectious Diseases (K08 AI136660). RSH was supported by NIH grants AI114852 and AG047773. JRG was supported by an NIH grant T32CA009140 and a Cancer Research Institute-Mark Foundation Fellowship. This work was supported by NIH grants (AI105343, AI082630, AI112521, AI108545,) to EJW; EJW is supported by the Parker Institute for Cancer Immunotherapy which supports the cancer immunology program at UPenn.

## AUTHOR CONTRIBUTIONS

RPS and EJW conceived the project. RPS, EJW, LAV, RSH, JRG, OK, JEW, and AEB helped in the design and interpretation of experiments. RPS performed the experiments with assistance from JEW, AEB, SM, and JRG. Bulk and single-cell RNAseq bioinformatics was performed by RPS, SM, and JRG. BCR sequencing and analysis was performed by WM, ETLP, and RPS. RPS, LAV, and EJW wrote the manuscript.

## STAR METHODS

### CONTACT FOR REAGENT AND RESOURCE SHARING

Further information and requests for reagents and resources should be directed to and will be fulfilled by the Lead Contact, E. John Wherry. Plasmid for expression of recombinant LCMV-NP was obtained from D. Pinschewer (University of Basel, Basel, Switzerland) and was originally developed by H. Pircher (University of Freiberg, Freiberg, Germany).

### EXPERIMENT MODEL AND SUBJECT DETAILS

#### Mice

6-8 week old C57BL/6 Ly5.2CR (CD45.1) or C57BL/6 (CD45.2) mice were purchased from NCI. Both male and female mice were used for these studies in accordance with NIH guidelines. Mice were maintained in a specific-pathogen-free facility at the University of Pennsylvania. All mice were used in accordance with Institutional Animal Care and Use Committee guidelines for the University of Pennsylvania.

## METHOD DETAILS

### Infections and antibody treatments

Mice were infected intraperitoneally (i.p.) with 2*10^5^ plaque-forming units (PFU) of LCMV Armstrong (Armstrong; acute) or intravenously (i.v.) with 4*10^6^ PFU LCMV clone 13 (clone 13; chronic). LCMV strains were propagated and titers were determined as previously described (Odorizzi et al., 2015; Wherry et al., 2003). For BCR inhibition experiments, ibrutinib (MedChem Express) was dissolved in CAPTEX355 (Abitec) at 1.25 mg/ml and mice were treated with 250ug of ibrutnib per dose as previously described (Mueller et al., 2015; Yang et al., 2016). For PE immunization experiments, recombinant phycoerythrin (PE; Prozyme) was diluted to 1mg/ml and 100ug was mixed with either LCMV Armstrong or LCMV clone 13 prior to infection. For repeated immunization experiments, mice were injected with either 200ul of PBS or 200ug of PE (Prozyme) i.p. at days 3, 6, 9, and 12 p.i.

### LCMV-NP protein production

Expression plasmids encoding a 6x His tagged version of the LCMV nucleoprotein (NP) (nucleoprotein sequences from LCMV Armstrong and LCMV clone 13 have 100% homology) were a kind gift from D. Pinschewer (University of Basel, Basel, Switzerland) and originally developed by H. Pircher (University of Freiberg, Germany) and was used as previously described (Fallet et al., 2016; Schweier et al., 2019; Sommerstein et al., 2015). Briefly, plasmid was transformed into Rosetta™ 2(DE3) competent cells (Novagen, Millipore Sigma). Protein expression was induced with 0.5M IPTG (Sigma) for 24 hours. Bacteria were harvested after induction, lysed and sonicated. The soluble fraction was purified over a HiTrap TALON crude column (GE Healthcare) on a GE AKTAprime plus FPLC system (GE Healthcare). Purified protein was desalted using PD-10 columns (GE Healthcare) and concentrated prior to use.

### Single-cell suspension preparation and LCMV-specific B cell staining and enrichment

Single-cell suspensions were obtained from spleens by homogenizing tissues against a 70um cell strainer. Cells were washed through a cell strainer and red blood cells were lysed in ACK lysis buffer (Thermo Fisher Scientific). Cells were resuspended in MACS buffer (PBS with 2mM EDTA and 1% fetal bovine serum) prior to cell staining or B cell enrichment. To identify LCMV-specific B cells, purified LCMV-NP was fluorescently labeled with Alexa Fluor 647 or Alexa Fluor 488 using antibody labeling kits according to manufacturer’s instructions (Thermo Fisher Scientific). Single-cell suspensions were stained with LCMV-NP for 15 minutes at 37°C in the presence of Fc-block (anti-CD16/32, 24G2, BD) and washed twice with MACS buffer prior to staining with additional fluorescent antibody panels. For bulk and single-cell RNAseq experiments, B cell enrichment (negative selection) was performed using mouse pan-B cell isolation kits (Stem Cell Technologies) according to manufacturer’s instructions except incubation times were doubled. Following B cell enrichment, LCMV-NP staining was performed as described above. LCMV-NP-labeled cell suspensions were further enriched (positive selection) for LCMV-NP-specific B cells by incubating cells with anti-Cy5/AF647 microbeads (Miltenyi Biotec) followed by purification over LS columns (Miltenyi Biotec).

### Flow cytometry and cell sorting

Single-cell suspensions were prepared and stained with fluorescently labeled LCMV-NP as described above. Cell suspensions were stained at 4°C for 30 minutes for surface protein staining. Intracellular staining was conducted at room temperature for 1 hour. Antibodies were purchased from Biolegend: CXCR4 (L276F12), CD83 (Michel-19), CD86 (GL-1), B220 (RA3-6B2), CD19 (1D3), IgD (11-26c.2a), CD138 (281-2), GL7 (GL7), CD38 (90), F4/80 (BM8), NK1.1 (PK136), IgM (RMM-1), Ki67 (16A8), CD69 (H1.2F3), Steptavidin-BV786; BD Biosciences: CXCR4 (2B11), CD83 (Michel-19), CD19, (1D3), CD138 (281-2), CD38 (90), CD4 (GK1.5), Blimp1 (5E7), Bcl6 (K112-91), CD31 (MEC 13.3), CCR6 (140706), Streptavidin BB780; eBioscience (Thermo Fisher Scientific): GL7 (GL7), CD4 (RM4-5), CD8 (53-6.7), F4/80 (BM8), NK1.1 (PK136); or from Southern Biotech: Ly6a (D7). Live cells were discriminated by staining with Live/Dead Aqua (Thermo Fisher Scientific) or Zombie Yellow (Biolegend). Intracellular and nuclear staining of transcription factors was performed using the FoxP3/Transcription Factor Staining Buffer Set (eBioscience, Thermo Fisher Scientific) according to manufacturer’s recommendations. Cell proliferation was measured by injecting mice i.p. with 3mg of 5-bromo-2’-deoxyuridine (BrdU) 2 hours prior to analysis. Fixation of cells for measurement of BrdU incorporation was done using a FIX & PERM Cell Permeabilization Kit (Thermo Fisher Scientific) with adjusted timings and BrdU was detected using an anti-BrdU antibody (MoBU-1, Thermo Fisher Scientifc). For cell sorting, samples were sorted directly into lysis buffer (RLT or RLT+, Qiagen) or into 10% FBS RPMI supplemented with 25mM HEPES. Flow cytometry data were acquired on a BD LSR II instrument or BD Symphony A3 instrument. Cell sorting was performed on a BD FACSAria enclosed within a laminar flow hood. Data were analyzed using FlowJo software (FlowJo, LLC).

### Immunofluorescent microscopy

Mouse spleen samples were acquired from LCMV Armstrong and LCMV-clone 13 infected mice at day 15 post-infection. Spleens were embedded in Optimal Cutting Temperature (OCT, Tissue-Tek, VWR) and snap frozen using a 2-methylbutane (Sigma) bath over dry ice. 7um sections on slides were stained as follows. Slides were desiccated at room temperature (RT) for 5-10 minutes prior to fixation in 4% freshly prepared paraformaldehyde for 10 minutes at RT. Following fixation, samples were rehydrated in PBS for 5 minutes followed by blocking for 1 hour in blocking buffer (1% bovine serum albumin (Sigma) and 0.3% Triton X-100 (Sigma)). Fluorescently-conjugated primary antibodies (IgD, clone 11-26c.2a, Biolegend and GL7, clone GL7, Thermo Fisher Scientific) were added and incubated for 2 hours at RT. Following incubation with primary antibodies, samples were wash 3x in PBS followed by coverslip mounting with ProLong Diamond with DAPI (Thermo Fisher Scientific).

Images were acquired on an inverted DMi8 widefield microscope (Leica) using a 40x objective (NA 0.95). At least 20 GCs were acquired per sample and GCs were sampled randomly across each tissue. Analysis of GC area was done in using the FIJI distribution (Schindelin et al., 2012) of ImageJ (Rueden et al., 2017) through manual annotation of GC boundaries followed by area determination.

### ELISA and antibody avidity ELISA

LCMV-specific IgG serum antibody titers were determined using standard ELISA protocols. Briefly, 96-well MaxiSorp plates (Nunc, Thermo Fisher Scientific) were coated with 300ng purified recombinant LCMV-NP per well overnight. For measurement of anti-LCMV-NP serum antibody concentration, serum samples were assayed in 5-fold dilutions starting from 1:100 dilution and plated in quadruplicate on each plate. LCMV-NP-specific IgG was detected using biotin-conjugated goat-anti-mouse IgG(H+L) (Southern Biotech) followed by incubation with streptavidin-HRP (Southern Biotech) and developed with tetramethylbenzidine (Thermo Fisher Scientific). Amount of antibody in each serum sample was calculated by fitting a standard curve generated using an anti-LCMV-NP monoclonal antibody (VL4, BioXcell).

Antibody avidity was measured by ELISA with modifications (Nawa, 1992; Pullen et al., 1986). For serum antibody avidity measurements, the anti-LCMV-NP ELISA was modified to include a wash step with either PBS, 1.5M sodium thiocyanate (NaSCN), 3M NaSCN, or 4.5M NaSCN for 15 minutes at room temperature after incubation of serum samples but before application of secondary goat-anti-mouse IgG(H+L) antibody. Avidity index was calculated as follows: Avidity index = (OD450_NaSCN_/OD450_PBS_)*100. Serum samples for serum avidity measurements were diluted to a concentration yielding OD450 of approximately 1 prior to incubation with NP-coated ELISA plates.

### B cell receptor sequencing (BCRseq)

All LCMV-NP-specific B cells were first sorted for yield then for purity from individual mice. Between 3,800 and 28,000 cells were obtained per sample. Samples were sorted into buffer RLT (Qiagen) + supplemented with β-mercaptoethanol (Sigma) and processed with an AllPrep DNA/RNA Micro Kit (Qiagen) according to manufacturer’s recommended protocol. Genomic DNA was sent to the Human Immunology Core at the University of Pennsylvania for mouse antibody heavy chain library construction, sequencing and data analysis. Primers used were adapted from Wang et. al. (Wang et al., 2000) at the beginning of the FW1 region of VH and were modified to include adaptor sequences for the Illumina NexteraXT kit (Illumina). Primer sequences used are below:

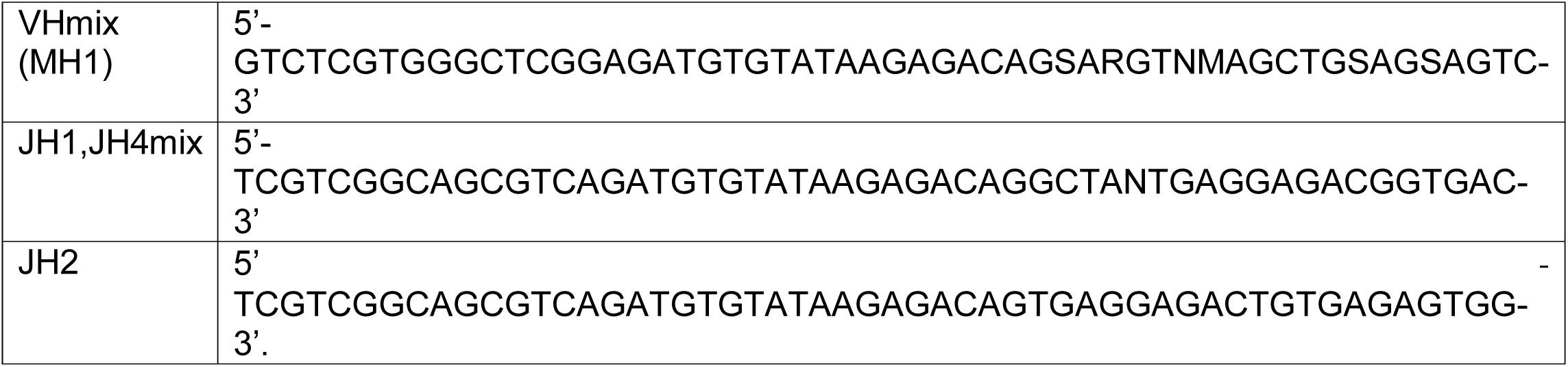

The mouse IgH library was generated with one VH primer and a cocktail of JH1,2,4 primers. The VH and JH primer mixes were used at 0.6 µM in a reaction volume of 25 µL using a Multiplex PCR kit (Qiagen, Valencia, CA, Cat. No. 158388). Amplification conditions for the PCR were: primary denaturation at 95°C for 10 minutes, cycling at 95°C 45s, 60°C 45s, and 72°C for 90s for 35 cycles, and a final extension step at 72°C for 10 minutes.

Amplicons were purified using the Agencourt AMPure XP beads system (Beckman Coulter, Inc., Indianapolis, IN), and second-round PCRs were performed as described (Meng et al., 2017) to add Illumina NexteraXT adaptors to the IgH library. Final sequencing libraries were quantified by Qubit Fluorometric Quantitation (Thermo Fisher Scientific, Grand Island, NY) and loaded onto an Illumina MiSeq instrument and sequenced using 2×300 bp paired end kits (Illumina MiSeq Reagent Kit v3, 600 cycle, Illumina Inc., San Diego, Cat. No. MS-102-3003).

Raw sequence data (FASTQ files) were processed through pRESTO v0.5.13 (Vander Heiden et al., 2014) and any sequence with a 10 bp section that had an average Phred quality score lower than 30 was removed, along with any sequence that was shorter than 100 bases. Germline V and J genes were identified by aligning pre-processed data to the murine IgH germline repertoire using the IMGT/HighV-QUEST tool hosted by the International Immunogenetics Information Systems (IMGT, imgt.org) (Alamyar et al., 2012; Lefranc et al., 2015; Li et al., 2013). Aligned reads were further processed with Change-O v0.4.6 (Gupta et al., 2015). The Change-O processing pipeline included filtering non-functional sequences and clustering sequences into clonal lineages. Data were then filtered by calculating the mean frequency (copy number) of all clones and selecting for clonal lineages above 50% of the mean copy number frequency. The frequency of mutation from germline, replacement to silent ratios (R/S ratio), and selection pressure (BASELINE, (Yaari et al., 2012)) were calculated using SHazaM v0.2.1 (Gupta et al., 2015).

### Bulk RNAseq sample preparation

To assess the kinetic transcriptional differences between LCMV-specific B cell subsets responding to LCMV Armstrong or LCMV-clone 13 infection, LCMV-specific GC B cells (CD38-GL7+), memory B cells (CD38+GL7-), and plasma cells (CD138+) were sorted from mice infected with LCMV Armstrong or LCMV clone 13 at days 7, 15, and 45 post-infection. All B cell subsets were first gated as LCMV-specific IgD-B220+Dump-prior to subset-specific gating. Dump gate included CD4, CD8, NK1.1, and F4/80. Naïve B cells were sorted from uninfected mice. Samples were sorted first for yield then for purity. 5-10 spleens were pooled for each of the 3 replicates sequenced at each timepoint. Prior to sorting, samples were enriched for LCMV-specific B cells and stained with surface antibodies as previously described. Sorted samples were stored in Buffer RLT supplemented with β-mercaptoethanol until RNA isolation. RNA isolation was performed with a RNeasy Mico Kit (Qiagen) per the manufacturer’s instructions. Library preparation was performed using a SMART-Seq v4 Ultra Low Input RNA kit (Takara) and Nextera XT DNA Library Preparation Kit (Illumina) per the manufacturer’s instructions. Quality of extracted RNA and prepared libraries were assessed using a TapeStation 2200 instrument (Agilent). Libraries were quantified using a KAPA Library Quantification Kit (Kapa Biosystems) prior to sequencing on an Illumina NextSeq 550 instrument (150bp, paired-end, 300 cycle high output flowcells).

### Bulk RNAseq data processing and analysis

FASTQ files were aligned using STAR 2.5.2a (Dobin et al., 2013) to the GRCm38-p5 murine reference genome (GENCODE, (Frankish et al., 2019)). Unique mapped reads were normalized using PORT v0.8.4-beta gene-based normalization (https://github.com/itmat/Normalization). Following PORT normalization, all downstream processing was conducted using R (R Core Team, 2019). Differential gene expression analysis was performed using Limma (Ritchie et al., 2015). Limma-voom was used to identify significantly different genes between groups using p value < 0.05. Ig genes were removed from each sample prior to differential gene expression analysis. Optimal K values for K-means clustering was calculated using the consensus K values calculated using the Elbow, Silhouette, and Gap Statistic methods of the NbClust package (Charrad et al., 2015). Heatmaps depict all differentially expressed genes (p value < 0.05 and log2 fold change > 2) within LCMV-specific B cell subsets between samples isolated from LCMV Armstrong and LCMV-clone 13 infected mice at days 7, 15, and 45 post-infection. Z-score values for genes in each cluster were calculated using the pheatmap package v1.0.12 (Kolde, 2019), averaged for each timepoint and infection, then plotted over time. Cluster Gene Ontology (GO) Biological Process term enrichment was performed using Metascape v3.0 (Zhou et al., 2019) with a minimum overlap of 3, p value cutoff of 0.01, and a minimum enrichment of 1.5. Gene set enrichment analysis (GSEA) was performed for gene sets of interest (Liberzon et al., 2015; Subramanian et al., 2005). Hallmark gene sets were obtained from MSigDB (Liberzon et al., 2015). GC B cell signature and spleen plasma cell signatures in were created using data deposited in the Gene Expression Omnibus (GEO, GSE60927, (Shi et al., 2015)). Briefly, gene signatures were generated by determining the top 200 differentially expressed genes between follicular B cells and subsets of interest as described above. Memory B cell signatures (Bhattacharya et al., 2007) and LZ and DZ GC B cell signatures (Victora et al., 2010) were previously published. Plots were made using ggplot2 (Wickham et al., 2019), the viridis package (Garnier, 2018), and the tidyverse package (Wickham, 2017).

### Single-cell RNAseq sample preparation

LCMV-specific GC B cells (CD38-GL7+) were sorted from mice infected with LCMV Armstrong or LCMV clone 13 at day 15 post-infection. Splenocytes from 5 mice per infection were pooled, enriched for LCMV-specific B cell and stained with surface antibodies as previously described. Samples were gated as LCMV-specific IgD-B220+Dump-prior to GC B cell gates. Dump stain included CD4, CD8, NK1.1, and F4/80. Cells were sorted for yield then for purity. Samples were counted using a Countess II Automated Cell Counter (Thermo Fisher Scientific) prior to library preparation. For library preparation, 5000 cells from each sample were loaded into a Chromium Controller (10x Genomics), emulsions were created, and libraries were constructed according to the manufacturer’s recommend protocols for the Chromium Single-cell 3’ v2 kit (10x Genomics). Samples were pooled and quantified using a KAPA Library Quantification Kit (Kapa Biosystems) prior to sequencing on an Illumina NextSeq 550 instrument (150 cycle, high output flow cell).

### Single-cell RNAseq data processing and analysis

BCL files from sequences samples were demultiplexed, aligned to the mouse mm10 genome, filtered, and UMI counted using Cell Ranger Software v2.1 (10x Genomics). 3,270 cells were recovered from LCMV Armstrong library with 88,576 mean reads per cell and 2,803 median genes detected per cell. 3,374 cells were recovered from LCMV clone 13 library with 83,259 mean reads per cell and 2,328 median genes detected per cell. Downstream analysis was performed using Seurat v2.3.1 (Butler et al., 2018). Complete script reproducing all analyses from raw data will be available in a Github repository upon publication of this data. Briefly, individual library outputs from the Cell Ranger pipeline were loaded, filtered to remove cells with > 5% mitochondrial reads and > 6000 unique genes detected. Filtered data from individual libraries were imputed using SAVER v0.3.0 (Huang et al., 2018) with default settings. Imputed count matrices were used for subsequent analysis in Seurat. Variable genes for each library were determined by calculating gene-gene correlations using Spearman distance, scoring gene-gene correlations, and taking the top 1500 highest correlated genes as described previously (https://hemberg-lab.github.io/scRNA.seq.course, section 8.3.2 Correlated Expression). Regression was performed to remove variation due to the number of genes detected or the percent of mitochondrial reads. Common correlation analysis (CCA) was performed to identify common sources of variation LCMV Armstrong and LCMV clone 13 prior to dataset merging using the first 12 CCA dimensions. Data was then clustered using the first 12 CCA dimensions and a resolution parameter of 0.6. Differential gene expression was conducted using the Wilcoxon rank sum test with a threshold of 0.25.

Single-cell enrichment scores for **Figure 4B, 4C, S4C,** and **S4D** were calculated using the AddModuleScore function in the Seurat package. Previously published gene sets were used to calculate LZ and DZ (Victora et al., 2010), PC, MBC vs. GC, and MBC vs PC single-cell enrichment scores (Bhattacharya et al., 2007; Shi et al., 2015). Cluster averaged single-cell enrichment scores for each gene signature of interest were used to define GC zone and differentiation state. Cell cycle scores were calculated using the CellCycleScoring function in the Seurat package using previously defined gene signatures for each cell cycle stage (Tirosh et al., 2016). GO term enrichment was conducted using Metascape v3.0 (Zhou et al., 2019) with a minimum overlap of 3, p value cutoff of 0.01, and a minimum enrichment of 1.5..

Pseudotime analysis of LZ GC B cells was conducted using Monocle 2 v2.10.1 (Qiu et al., 2017b, 2017a). CCA dimensions were imported from Seurat and used for pseudotime calculations. Pseudotime trajectory start was inferred using cell cycle staging and cluster function. Trajectories were rooted to start in cluster 4. Single-cell enrichment scores for gene sets of interest were calculated using GSVA (Hänzelmann et al., 2013). Preprocessed, filtered, normalized, and log-transformed scRNAseq expression data was used as input to GSVA treating each cell as a unique sample. A Gaussian kernel distribution was used for the GSVA calculation as input data was previously log-transformed. Single-cell GSVA scores were plotted as a function of pseudotime to obtain the graphs in **Figures 6A, 6E**, and **S6B-G**.

## QUANTIFICATION AND STATISTICAL ANALYSIS

For each figure, the n, representing the number of mice per group, and the number of replicates for each experiment are indicated in the figure legends. In all figures, error bars represent the ± SEM. Paired Student’s t tests (two-tailed), Mann Whitney U tests, and one-way ANOVAs with Tukey’s multiple comparison test were used for statistical analysis when appropriate and the exact test used for each statistical analysis used is indicated in the figure legends. Individual points represent one mouse unless indicated. For longitudinal data, in Figure 1, S1, ELISA data in Figure 2, and S2 each point plotted is the mean and Student’s t test was used to test differences between acute and chronic infection at each time point. Each point in Figure S1D represents one GC from n = 2 mice per group. Data in Figure 2E are pooled from 5 individual BCR repertoires from each infection group. Data in Figure S5C and S5D are averaged single-cell enrichment scores for the indicated gene signatures from cells comprising each cluster. Asterisks indicate statistical significance (* p < 0.05; ** p < 0.01, *** p < 0.001). Statistical analyses were conducted using Prism software v8.2.1 (GraphPad) or R software v3.6 (The R Foundation for Statistical Computing).

## SUPPLEMENTAL INFORMATION

**Supplementary Figure 1: LCMV-specific B cell responses are diminished during chronic infection. Related to Figure 1.**

**(A)** Gating strategy for identifying LCMV-specific B cells and LCMV-specific GC B cells, MBC, and PC. Cells are first gated for live lymphocyte singlets before B cell gating. **(B)** Enumeration of LCMV-specific B cell response in LCMV Armstrong (acute) and LCMV clone 13 (chronic) at days 7, 10, 15, 21, 35, and 110 post-infection. **(C)** Expression of Tbet in CD38-GL7-CD138-B cells as gated in A. **(D)** 40x images of GCs in LCMV Armstrong and LCMV clone 13 infected mice at day 15 post-infection. GC size was quantitated for two individual spleens per infection by staining IgD (red) and GL7 (blue). At least 20 GCs per sample were enumerated. Size was manually determined by marking border between IgD+ cells in the follicular mantle zone and GL7+ cells within the GC as depicted by yellow dashed line in representative images. **(D-E)** Enumeration of LCMV-specific splenic PC **(D)**, GC B cell **(E)**, and MBC **(F)** in LCMV Armstrong and LCMV clone 13 at days 7, 10, 15, 21, 35, and 110 post-infection. Each data point in A and C-D shows mean ± SEM with 3-5 mice per time point from two independent experiments. Data points in D show mean ± SEM from 10 mice pooled from two independent experiments. *p<0.05, **p<0.01, ***p<0.001, Student’s t-test

**Supplementary Figure 2: Chronic infection impairs affinity maturation but BCR repertoire usage is largely unaffected. Related to Figure 2.**

**(A)** Anti-LCMV-NP serum avidity index in response to NaSCN treatment in LCMV Armstrong (acute) and LCMV clone 13 (chronic) infected mice at days 7, 10, 15, 21, 35, and 110 post-infection. **(B)** Frequency of heavy chain BCR repertoire represented by the 20 most abundant clonal families from LCMV Armstrong and LCMV clone 13 infection mice at day 45 post-infection. **(C)** Heavy chain VH1 gene usage in LCMV-specific B cell BCR repertoires from LCMV Armstrong and LCMV clone 13 infection mice at day 45 post-infection. Each point represents one mouse. Data representative of two independent experiments. **(D)** Plot of BCR family clonal abundance by average mutation frequency. **(E)** Frequency of heavy chain mutations in broken down by heavy chain region. Complementarity determining region (CDR) and framework (FWR). **(F)** Frequency of replacement and silent mutations in BCR repertoires analyzed. Data are pooled from IgH BCR repertoires of 5 individual mice per infection. ***p<0.001, Student’s t-test.

**Supplementary Figure 3: Transcriptional regulation of LCMV-specific GC B cells, PC, and MB is temporally dynamic. Related to Figure 3.**

**(A)** Purification and FACS sorting strategy used for Figures 3, 4 and 5 (BCR sequencing, bulk RNAseq, and scRNA-seq). Strategy is as follows: 1) Bulk splenocytes are first negatively selected for pan-B cells using magnetic depletion, 2) Enriched B cells are stained with AF647-conjugated LCMV-NP and magnetic positive selection is done using anti-AF647 microbeads, 3) Enriched LCMV-specific B cells are then stained for FACS sorting and sorted for yield, 4) Yield-sorted LCMV-specific B cells are finally sorted for purity and populations of interest. Representative FACS plots depict B cell and LCMV-specific B cell frequencies following each step in the purification process. **(B-D)** Heatmap depicting all differentially expressed (p<0.05 and >2 log fold-change) genes between LCMV-specific GC B cells **(B)**, plasma cells **(C)**, and memory B cells **(D)** from LCMV Armstrong (acute) and LCMV clone 13 (chronic) infected mice at days 7, 15, or 45 post-infection. Selected genes are shown. Heatmaps were k-means clustered and average z-scores for each cluster were plotted over time for each infection. **(E)** GSEA plot of Hallmark MTORC1 signaling, Hallmark MYC Targets V1, and Hallmark MYC Targets V2 gene signatures in LCMV-specific GC B cells from LCMV Armstrong and LCMV clone 13 infected mice at day 15 post-infection.

**Supplementary Figure 4: Unsupervised clustering of LCMV-specific GC B cells identifies clusters with unique biology and known GC B cell function. Related to Figure 4.**

**(A)** Heatmap of LCMV-specific GC B cell cluster-defining genes. Heatmap depicts top 10 most differentially expressed genes per cluster. **(B)** Top enriched GO terms for each identified cluster in LCMV-specific GC B cells scRNAseq data. **(C)** Averaged single-cell enrichment scores for the LZ and DZ gene signatures in each cluster. **(D)** Averaged single-cell enrichment scores for genes upregulated in PC, genes upregulated in MBC versus GC B cells, and gene upregulated in MBC vs PC for each cluster. **(E)** Frequency of cells in each cluster in the indicated cell cycle stage.

**Supplementary Figure 5: Pseudotime analysis identifies GC B cell fate decision points within the LZ and correlated gene expression signatures. Related to Figures 5 and 6.**

**(A)** Pseudotime analysis of light zone GC B cell populations identified using scRNAseq. Pseudotime is rooted in cluster 4: T cell and FDC interactions. Plots depict pseudotime, cell cycle stage, cluster identity, and distinct pseudotime states of LCMV-specific LZ GC B cells from both LCMV Armstrong (acute) and LCMV clone 13 (chronic) infection. **(B)** Single-cell GSVA enrichment scores for Basso CD40 signaling genes UP geneset plotted as a function of pseudotime. **(C)** Single-cell GSVA enrichment scores for Hallmark PI3K-MTOR-AKT signaling geneset plotted as a function of pseudotime. **(D)** Single-cell GSVA enrichment scores for Hallmark MTORC1 signaling geneset plotted as a function of pseudotime. **(E)** Single-cell GSVA enrichment scores for Hallmark MYC targets V2 geneset plotted as a function of pseudotime. **(F)** Single-cell GSVA enrichment scores for Hallmark Response to Interferon Alpha geneset plotted as a function of pseudotime. **(G)** Single-cell GSVA enrichment scores for Hallmark Response to Interferon Gamma geneset plotted as a function of pseudotime.

**Supplementary Figure 6: BCR signaling pathways but not inhibitory receptor expression is altered during chronic viral infection. Related to Figure 6.**

**(A)** Expression of B cell inhibitory receptors and phosphatases in LCMV-specific GC B cells within cluster 4 (T cell FDC interactions) from both LCMV Armstrong (acute) and LCMV clone 13 (chronic) infected mice at day 15 post-infection. **(B)** Expression of selected genes downstream of the BCR in LCMV-specific GC B cells in cluster 4 (T cell FDC interactions). ***p<0.001 Wicoxon Rank Sum Test.

