## Supplemental Figures for "Chronic viral infection promotes early germinal center exit of B cells and impaired antibody development"

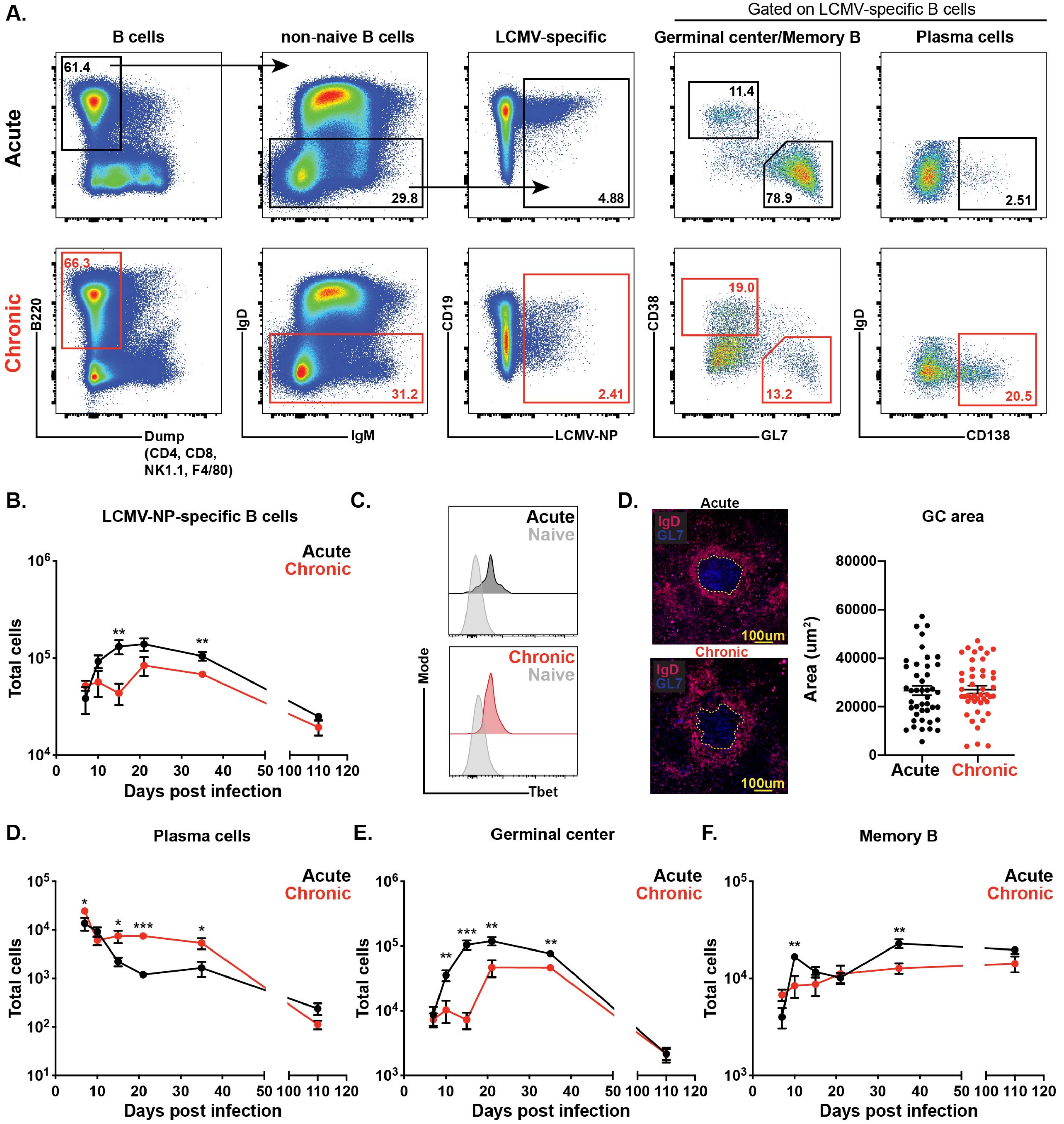
A.
Gated on LCMV-specific B cells
B cells
non-naive B cells
LCMV-specific
Germinal center/Memory B
Plasma cells
Acute
Chronic
61.4
29.8
4.88
11.4
78.9
2.51
66.3
31.2
2.41
19.0
13.2
20.5
B220
Dump
(CD4, CD8,
NK1.1, F4/80)
IgD
IgM
CD19
LCMV-NP
CD38
GL7
IgD
CD138
B.
LCMV-NP-specific B cells
Acute
Chronic
Total cells
Days post infection
C.
Acute
Naive
Chronic
Naive
Mode
Tbet
D.
Acute
IgD
GL7
100um
Chronic
GC area
Area (um²)
Acute
Chronic
D.
Plasma cells
E.
Germinal center
F.
Memory B
Total cells
Days post infection

**Supplementary Figure 1: LCMV-specific B cell responses are diminished during chronic infection. Related to Figure 1.**

**(A)** Gating strategy for identifying LCMV-specific B cells and LCMV-specific GC B cells, MBC, and PC. Cells are first gated for live lymphocyte singlets before B cell gating. **(B)** Enumeration of LCMV-specific B cell response in LCMV Armstrong (acute) and LCMV clone 13 (chronic) at days 7, 10, 15, 21, 35, and 110 post-infection. **(C)** Expression of Tbet in CD38-GL7-CD138- B cells as gated in A. **(D)** 40x images of GCs in LCMV Armstrong and LCMV clone 13 infected mice at day 15 post-infection. GC size was quantitated for two individual spleens per infection by staining IgD (red) and GL7 (blue). At least 20 GCs per sample were enumerated. Size was manually determined by marking border between IgD+ cells in the follicular mantle zone and GL7+ cells within the GC as depicted by yellow dashed line in representative images. **(D-E)** Enumeration of LCMV-specific splenic PC **(D)**, GC B cell **(E)**, and MBC **(F)** in LCMV Armstrong and LCMV clone 13 at days 7, 10, 15, 21, 35, and 110 post-infection. Each data point in A and C-D shows mean ± SEM with 3-5 mice per time point from two independent experiments. Data points in D show mean ± SEM from 10 mice pooled from two independent experiments. *p<0.05, **p<0.01, ***p<0.001, Student's t-test

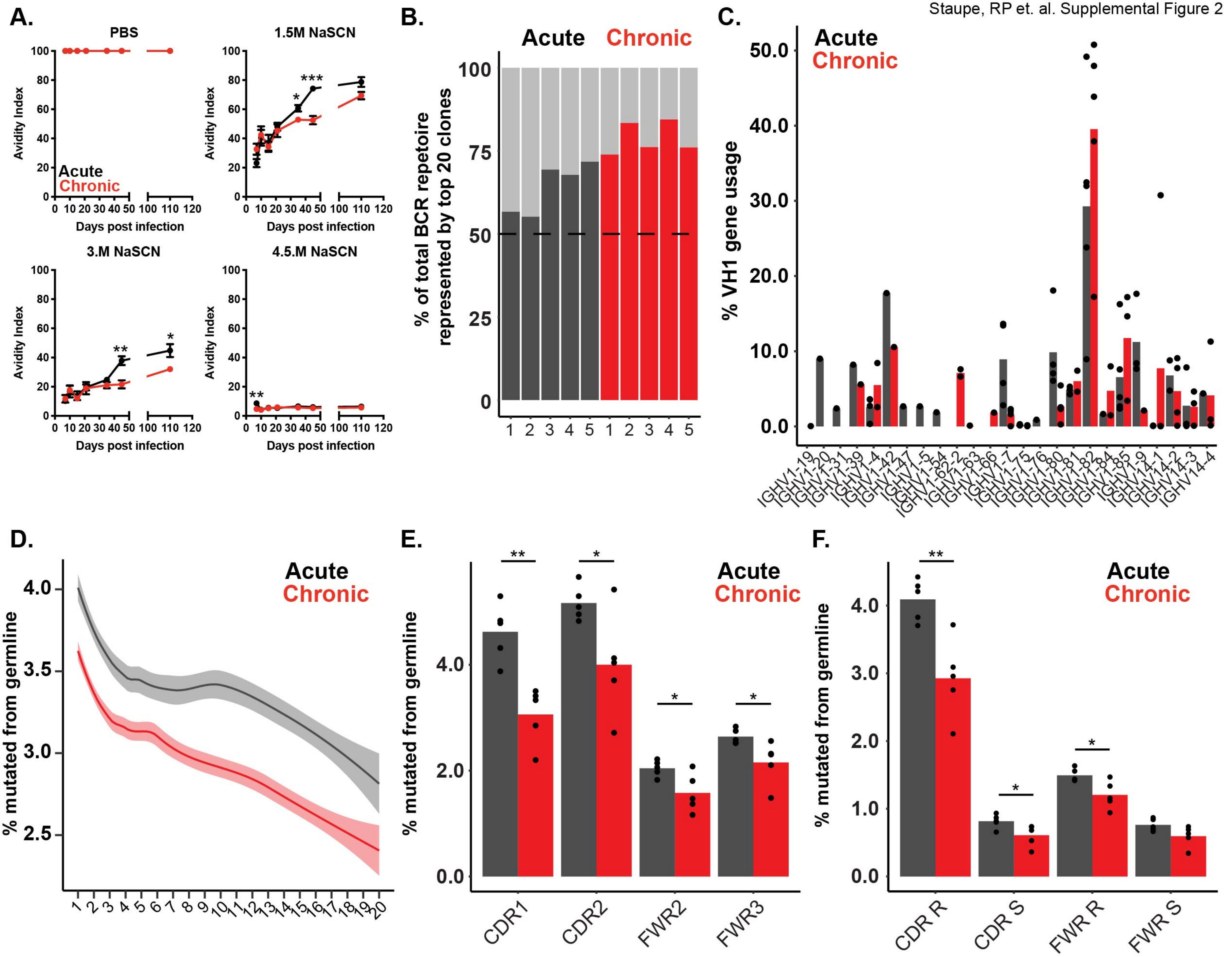
A.
PBS
1.5M NaSCN
3.M NaSCN
4.5.M NaSCN
Avidity Index
Days post infection
Acute
Chronic
B.
Acute Chronic
% of total BCR repetoire represented by top 20 clones
C.
Acute
Chronic
% VH1 gene usage
IGHV1-19
IGHV1-20
IGHV1-31
IGHV1-39
IGHV1-4
IGHV1-42
IGHV1-47
IGHV1-5
IGHV1-54
IGHV1-62-2
IGHV1-63
IGHV1-66
IGHV1-7
IGHV1-75
IGHV1-76
IGHV1-80
IGHV1-81
IGHV1-82
IGHV1-84
IGHV1-85
IGHV1-9
IGHV14-1
IGHV14-2
IGHV14-3
IGHV14-4
D.
Acute
Chronic
% mutated from germline
E.
Acute
Chronic
% mutated from germline
CDR1
CDR2
FWR2
FWR3
F.
Acute
Chronic
% mutated from germline
CDR R
CDR S
FWR R
FWR S

**Supplementary Figure 2: Chronic infection impairs affinity maturation but BCR repertoire usage is largely unaffected. Related to Figure 2.**

**(A)** Anti-LCMV-NP serum avidity index in response to NaSCN treatment in LCMV Armstrong (acute) and LCMV clone 13 (chronic) infected mice at days 7, 10, 15, 21, 35, and 110 post-infection. **(B)** Frequency of heavy chain BCR repertoire represented by the 20 most abundant clonal families from LCMV Armstrong and LCMV clone 13 infection mice at day 45 post-infection. **(C)** Heavy chain VH1 gene usage in LCMV-specific B cell BCR repertoires from LCMV Armstrong and LCMV clone 13 infection mice at day 45 post-infection. Each point represents one mouse. Data representative of two independent experiments. **(D)** Plot of BCR family clonal abundance by average mutation frequency. **(E)** Frequency of heavy chain mutations in broken down by heavy chain region. Complementarity determining region (CDR) and framework (FWR). **(F)** Frequency of replacement and silent mutations in BCR repertoires analyzed. Data are pooled from IgH BCR repertoires of 5 individual mice per infection. ***$p<0.001$, Student's t-test.

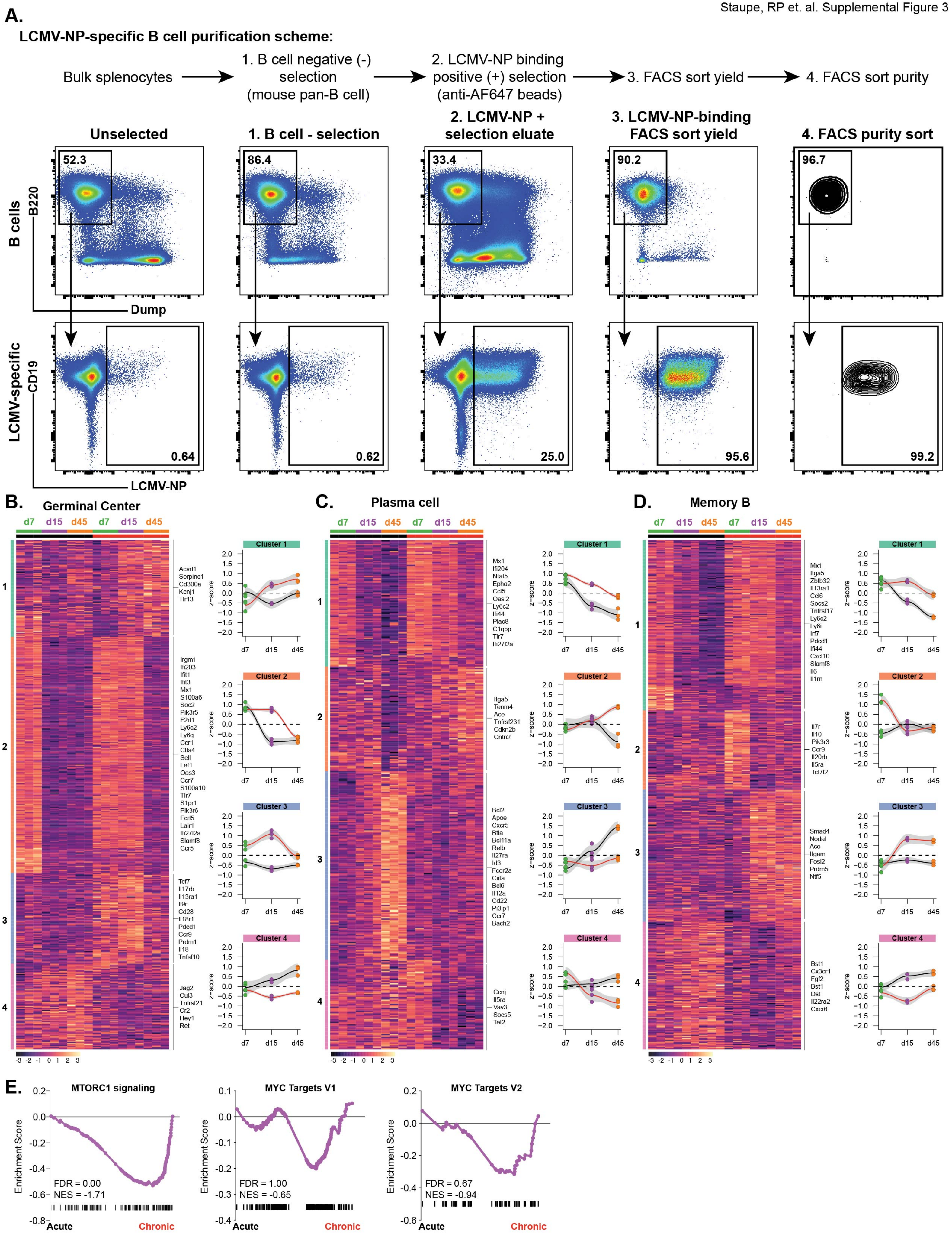
A.
LCMV-NP-specific B cell purification scheme:
Bulk splenocytes
1. B cell negative (-) selection (mouse pan-B cell)
2. LCMV-NP binding positive (+) selection (anti-AF647 beads)
3. FACS sort yield
4. FACS sort purity
Unselected
1. B cell - selection
2. LCMV-NP + selection eluate
3. LCMV-NP-binding FACS sort yield
4. FACS purity sort
52.3
86.4
33.4
90.2
96.7
B cells
B220
Dump
LCMV-specific
CD19
LCMV-NP
0.64
0.62
25.0
95.6
99.2
B.
Germinal Center
d7 d15 d45 d7 d15 d45
1
2
3
4
Acvrl1
Serpinc1
Cd300a
Kcnj1
Tlr13
Irgm1
Ifi203
Ifit1
Ifit3
Mx1
S100a6
Soc2
Pik3r5
F2rl1
Ly6c2
Ly6g
Ccr1
Ctla4
Sell
Lef1
Oas3
Ccr7
S100a10
Tlr7
S1pr1
Pik3r6
Fcrl5
Lair1
Ifi27l2a
Slamf8
Ccr5
Tcf7
Il17rb
Il13ra1
Il9r
Cd28
Il18r1
Pdcd1
Ccr9
Prdm1
Il18
Tnfsf10
Jag2
Cul3
Tnfrsf21
Cr2
Hey1
Ret
Cluster 1
Cluster 2
Cluster 3
Cluster 4
z-score
d7 d15 d45
-3 -2 -1 0 1 2 3
C.
Plasma cell
Mx1
Ifi204
Nfat5
Epha2
Ccl5
Oasl2
Ly6c2
Ifi44
Plac8
C1qbp
Tlr7
Ifi27l2a
Itga5
Tenm4
Ace
Tnfrsf231
Cdkn2b
Cntn2
Bcl2
Apoe
Cxcr5
Btla
Bcl11a
Relb
Il27ra
Id3
Fcer2a
Ciita
Bcl6
Il12a
Cd22
Pi3ip1
Ccr7
Bach2
Ccnj
Il5ra
Vav3
Socs5
Tet2
D.
Memory B
Mx1
Itga5
Zbtb32
Il13ra1
Ccl6
Socs2
Tnfrsf17
Ly6c2
Ly6i
Irf7
Pdcd1
Ifi44
Cxcl10
Slamf8
Il6
Il1m
Il7r
Il10
Pik3r3
Ccr9
Il20rb
Il5ra
Tcf7l2
Smad4
Nodal
Ace
Itgam
Fosl2
Prdm5
Ntf5
Bst1
Cx3cr1
Fgf2
Bst1
Dst
Il22ra2
Cxcr6
E.
MTORC1 signaling
FDR = 0.00
NES = -1.71
MYC Targets V1
FDR = 1.00
NES = -0.65
MYC Targets V2
FDR = 0.67
NES = -0.94
Enrichment Score
Acute
Chronic

**Supplementary Figure 3: Transcriptional regulation of LCMV-specific GC B cells, PC, and MB is temporally dynamic. Related to Figure 3.**

**(A)** Purification and FACS sorting strategy used for Figures 3, 4 and 5 (BCR sequencing, bulk RNAseq, and scRNA-seq). Strategy is as follows: 1) Bulk splenocytes are first negatively selected for pan-B cells using magnetic depletion, 2) Enriched B cells are stained with AF647-conjugated LCMV-NP and magnetic positive selection is done using anti-AF647 microbeads, 3) Enriched LCMV-specific B cells are then stained for FACS sorting and sorted for yield, 4) Yield-sorted LCMV-specific B cells are finally sorted for purity and populations of interest. Representative FACS plots depict B cell and LCMV-specific B cell frequencies following each step in the purification process. **(B-D)** Heatmap depicting all differentially expressed ($p<0.05$ and >2 log fold-change) genes between LCMV-specific GC B cells **(B)**, plasma cells **(C)**, and memory B cells **(D)** from LCMV Armstrong (acute) and LCMV clone 13 (chronic) infected mice at days 7, 15, or 45 post-infection. Selected genes are shown. Heatmaps were k-means clustered and average z-scores for each cluster were plotted over time for each infection. **(E)** GSEA plot of Hallmark MTORC1 signaling, Hallmark MYC Targets V1, and Hallmark MYC Targets V2 gene signatures in LCMV-specific GC B cells from LCMV Armstrong and LCMV clone 13 infected mice at day 15 post-infection.

Staupe, RP et. al. Supplemental Figure 4

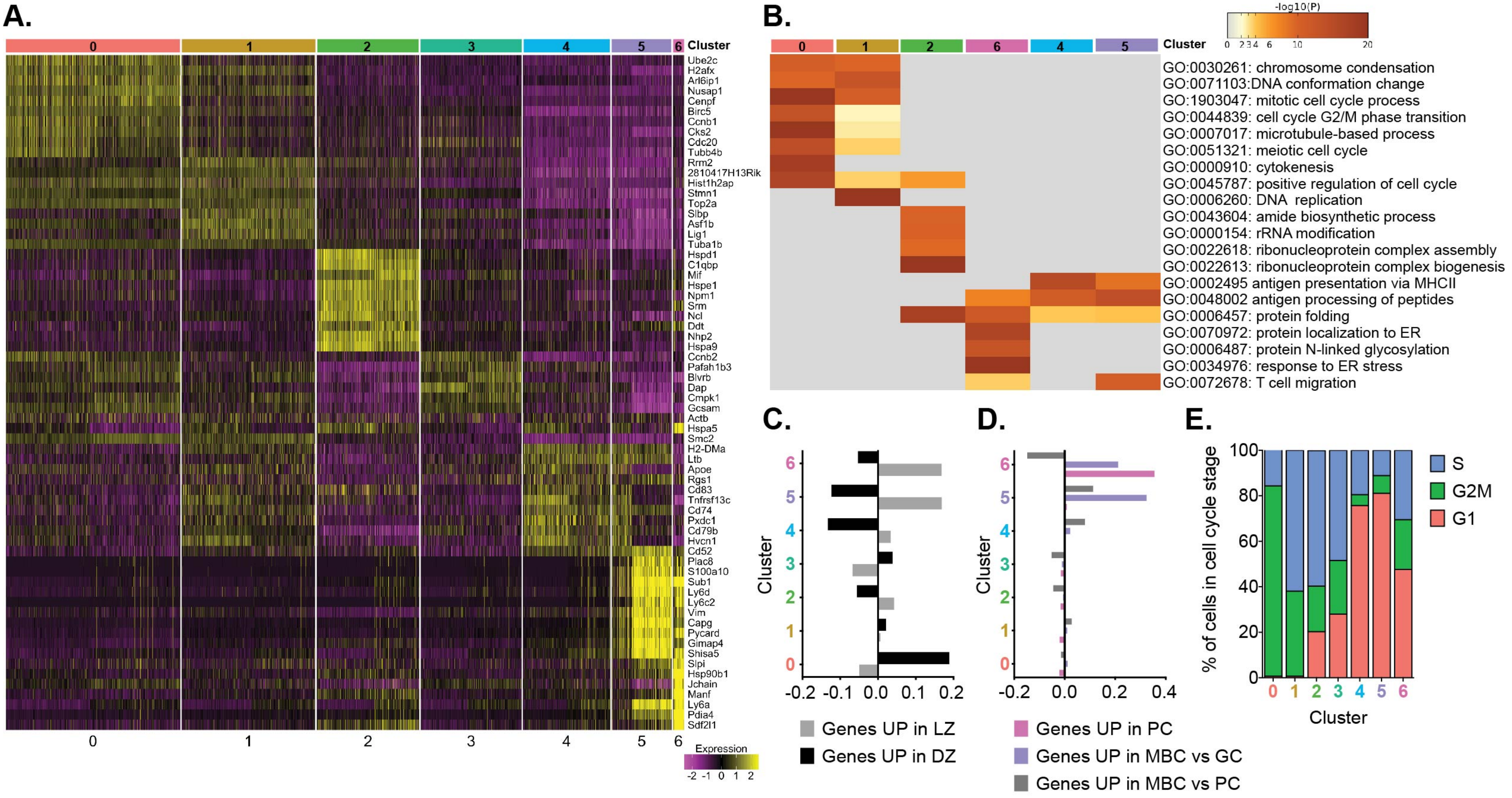

**Supplementary Figure 4: Unsupervised clustering of LCMV-specific GC B cells identifies clusters with unique biology and known GC B cell function. Related to Figure 4.**

**(A)** Heatmap of LCMV-specific GC B cell cluster-defining genes. Heatmap depicts top 10 most differentially expressed genes per cluster. **(B)** Top enriched GO terms for each identified cluster in LCMV-specific GC B cells scRNAseq data. **(C)** Averaged single-cell enrichment scores for the LZ and DZ gene signatures in each cluster. **(D)** Averaged single-cell enrichment scores for genes upregulated in PC, genes upregulated in MBC versus GC B cells, and gene upregulated in MBC vs PC for each cluster. **(E)** Frequency of cells in each cluster in the indicated cell cycle stage.

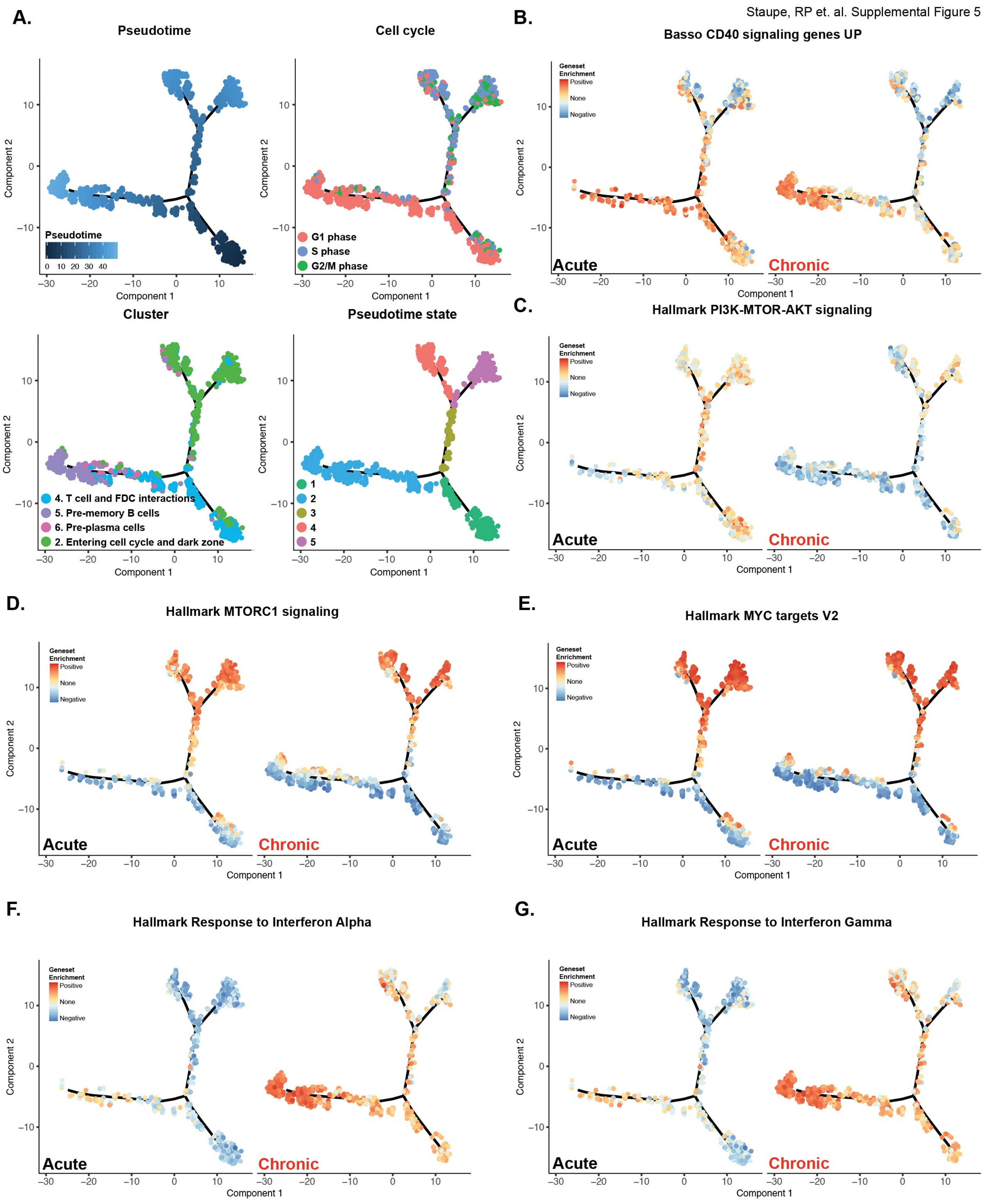
A.
Pseudotime
Cell cycle
G1 phase
S phase
G2/M phase
Cluster
4. T cell and FDC interactions
5. Pre-memory B cells
6. Pre-plasma cells
2. Entering cell cycle and dark zone
Pseudotime state
Component 1
Component 2
B.
Basso CD40 signaling genes UP
C.
Hallmark PI3K-MTOR-AKT signaling
D.
Hallmark MTORC1 signaling
E.
Hallmark MYC targets V2
F.
Hallmark Response to Interferon Alpha
G.
Hallmark Response to Interferon Gamma
Geneset Enrichment
Positive
None
Negative
Acute
Chronic

**Supplementary Figure 5: Pseudotime analysis identifies GC B cell fate decision points within the LZ and correlated gene expression signatures. Related to Figures 5 and 6.**

**(A)** Pseudotime analysis of light zone GC B cell populations identified using scRNAseq. Pseudotime is rooted in cluster 4: T cell and FDC interactions. Plots depict pseudotime, cell cycle stage, cluster identity, and distinct pseudotime states of LCMV-specific LZ GC B cells from both LCMV Armstrong (acute) and LCMV clone 13 (chronic) infection. **(B)** Single-cell GSVA enrichment scores for Basso CD40 signaling genes UP geneset plotted as a function of pseudotime. **(C)** Single-cell GSVA enrichment scores for Hallmark PI3K-MTOR-AKT signaling geneset plotted as a function of pseudotime. **(D)** Single-cell GSVA enrichment scores for Hallmark MTORC1 signaling geneset plotted as a function of pseudotime. **(E)** Single-cell GSVA enrichment scores for Hallmark MYC targets V2 geneset plotted as a function of pseudotime. **(F)** Single-cell GSVA enrichment scores for Hallmark Response to Interferon Alpha geneset plotted as a function of pseudotime. **(G)** Single-cell GSVA enrichment scores for Hallmark Response to Interferon Gamma geneset plotted as a function of pseudotime.

Staupe, RP et. al. Supplemental Figure 6

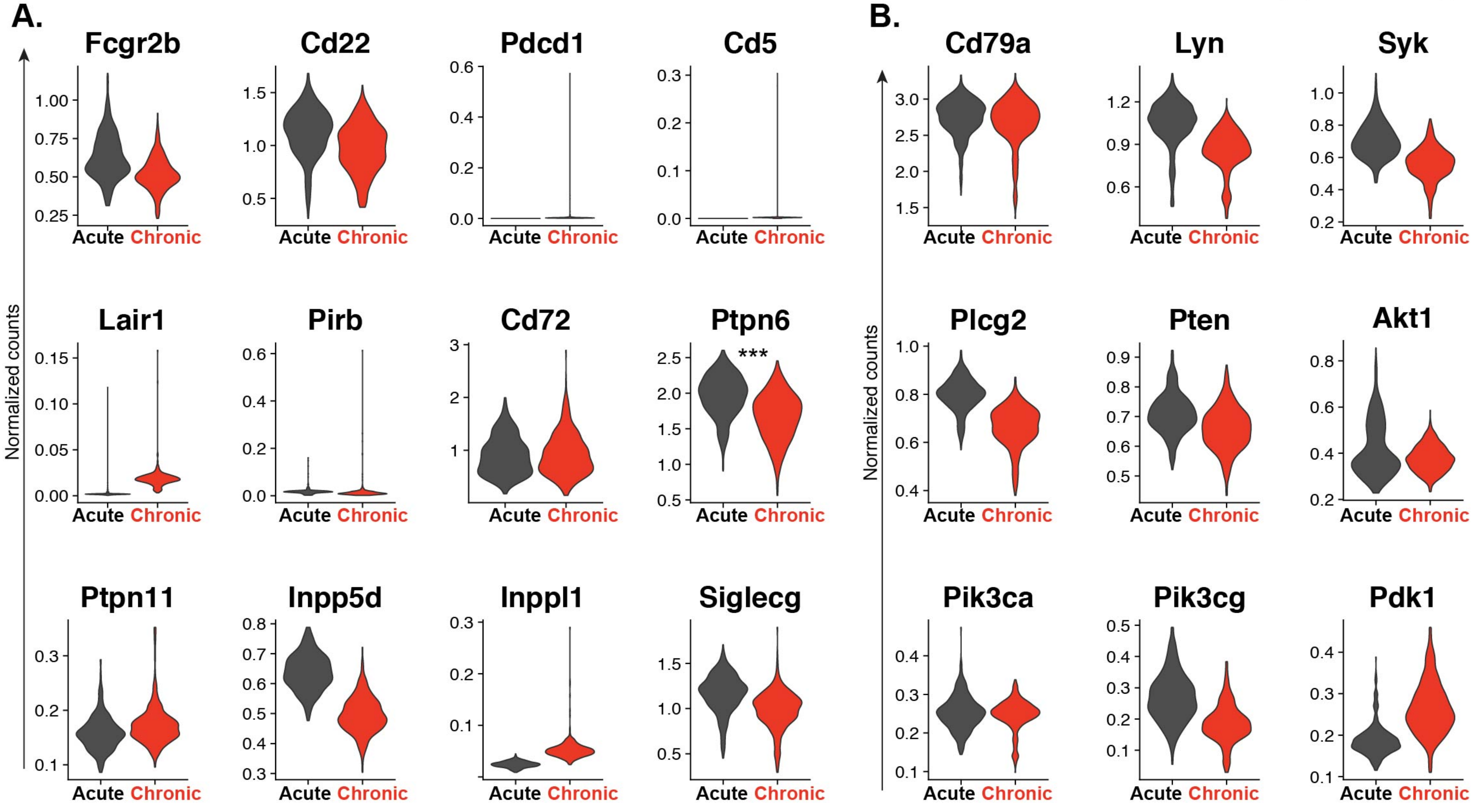

**Supplementary Figure 6: BCR signaling pathways but not inhibitory receptor expression is altered during chronic viral infection. Related to Figure 6.**

**(A)** Expression of B cell inhibitory receptors and phosphatases in LCMV-specific GC B cells within cluster 4 (T cell FDC interactions) from both LCMV Armstrong (acute) and LCMV clone 13 (chronic) infected mice at day 15 post-infection. **(B)** Expression of selected genes downstream of the BCR in LCMV-specific GC B cells in cluster 4 (T cell FDC interactions). ***p<0.001 Wicoxon Rank Sum Test.
